# Suppressing DNMT3a Alleviates the Intrinsic Epigenetic Barrier for Optic Nerve Regeneration and Restores Vision in Adult Mice

**DOI:** 10.1101/2023.11.17.567614

**Authors:** Wai Lydia Tai, Kin-Sang Cho, Emil Kriukov, Ajay Ashok, Xuejian Wang, Aboozar Monavarfeshani, Wenjun Yan, Yingqian Li, Timothy Guan, Joshua R. Sanes, Petr Baranov, Dong Feng Chen

**Affiliations:** Schepens Eye Research Institute of Mass Eye and Ear, Department of Ophthalmology, Harvard Medical School, Boston, MA, USA; School of Pharmacy, Weifang Medical University, Weifang, Shandong, China; Department of Cellular and Molecular Biology, Center for Brain Science, Harvard University, MA, USA

## Abstract

The limited regenerative potential of the optic nerve in adult mammals presents a major challenge for restoring vision after optic nerve trauma or disease. The mechanisms of this regenerative failure are not fully understood^1,2^. Here, through small-molecule and genetic screening for epigenetic modulators^3^, we identify DNA methyltransferase 3a (DNMT3a) as a potent inhibitor of axon regeneration in mouse and human retinal explants. Selective suppression of DNMT3a in retinal ganglion cells (RGCs) by gene targeting or delivery of shRNA leads to robust, full-length regeneration of RGC axons through the optic nerve and restoration of vision in adult mice after nerve crush injury. Genome-wide bisulfite and transcriptome profiling in combination with single nucleus RNA-sequencing of RGCs revealed selective DNA demethylation and reactivation of genetic programs supporting neuronal survival and axonal growth/regeneration by DNMT3a deficiency. This was accompanied by the suppression of gene networks associated with apoptosis and inflammation. Our results identify DNMT3a as the central orchestrator of an RGC-intrinsic mechanism that limits optic nerve regeneration. Suppressing DNMT3a expression in RGCs unlocks the epigenetic switch for optic nerve regeneration and presents a promising therapeutic avenue for effectively reversing vision loss resulted from optic nerve trauma or diseases.

## MAIN TEXT

Mature neurons in the central nervous system (CNS) of adult mammals regenerate poorly after injury. Tremendous efforts have been devoted to study the cellular and molecular origins of this regenerative failure, yet no treatment exists for successful stimulation of CNS regeneration in humans. Eye-to-brain circuits, consisting of retinal ganglion cells (RGCs) whose axons form the optic nerve and connect to the central target areas, is an attractive model system for investigation of factors that regulate CNS axon regeneration because of their accessibility and simple anatomy. However, despite decades of research, achieving functional regeneration of the optic nerve has proved to be an elusive goal. Optic nerve regeneration requires a complex sequence of steps controlled by intricate gene networks that mediate crucial processes such as RGC survival, axonal elongation and guidance, and reformation of synapsis and neural circuitry^1,4^. The current prevailing view is that functional regeneration of the optic nerve cannot be achieved by the activation of a single gene or transcription factor^1,2^, but rather requires modulation of multiple gene expression or programs.

Embryonic RGCs possess the innate ability to regenerate optic nerve fibers and readily grow their axons even when presented with a hostile or adult brain environment (growth-inhibitory)^5,6^. However, this ability of RGCs is lost perinatally^4-7^ at a time coinciding with dynamic changes in epigenetic factors during RGC maturation^8^. We hypothesized that RGC axon growth programs are switched off during maturation through an epigenetic silencing mechanism involving DNA methylation. Indeed, a recent study demonstrated promising results in reprogramming the epigenomes of adult RGCs through the ectopic expression of three transcription factors used to generate stem cells – Oct4, Sox2, and Klf4 (Yamanaka factors)^9^. This resulted in partial restoration of youthful DNA methylation patterns and significant axon regeneration following crush injury. To date, however, no known strategy fully restores optic nerve regeneration with visual recovery following nerve injury in adult mammals. We sought strategies to unleash and completely reverse the epigenetic blockade and restore the regenerative potential of the optic nerve in adult mammals. We show here that DNMT3a is pivotal for postnatal onset of optic nerve regenerative failure, and suppression of this single gene results in reprogramming of RGC transcriptomic landscape to enable unprecedented level of optic nerve regeneration and reversal of vision loss in adult mice.

### RGC axon growth enabled by *Dnmt3a* deficiency

Mouse RGCs lose their intrinsic ability to regenerate axons around embryonic day 18 (E18), one day before birth (postnatal day 0; P0)^5^. To assess epigenetic mechanisms responsible for this decline, we examined the expression patterns of DNA methyltransferases (DNMTs) in developing RGCs of mouse pups aged E16, P0, and P10, which is, before, at, and after the time that RGCs lose their ability to regenerate axons. Quantitative polymerase chain reaction (qPCR) of purified RGCs showed that the expression of *Dnmt3a* increased significantly from E16 to P10, which corresponded to the decline of RGCs’ axon regenerative capacity (**Fig. 1a**). Other DNMTs, including *Dnmt1* and *Dnmt3b*, did not exhibit this correlation (**Extended Data** Fig. 1a). Immunohistochemistry confirmed this result: expression of DNMT3a was absent in E16 RGCs or retina but intense in P10 retina (**Fig. 1b** and **Extended Data** Fig. 1b), which was especially pronounced in the RGCs as evident by colocalization with RBPMS, an RGC marker (**Fig. 1b**). Moreover, *Dnmt3a* levels were increased in the RGCs two days after optic nerve crush injury (ONC; **Fig. 1c**), while *Dnmt1* and *Dnmt3b* remained unchanged or downregulated (**Extended Data** Fig. 1c). These data suggest a correlation between DNMT3a upregulation and the decrease of optic nerve regenerative capacity in mice.

**Fig. 1.**
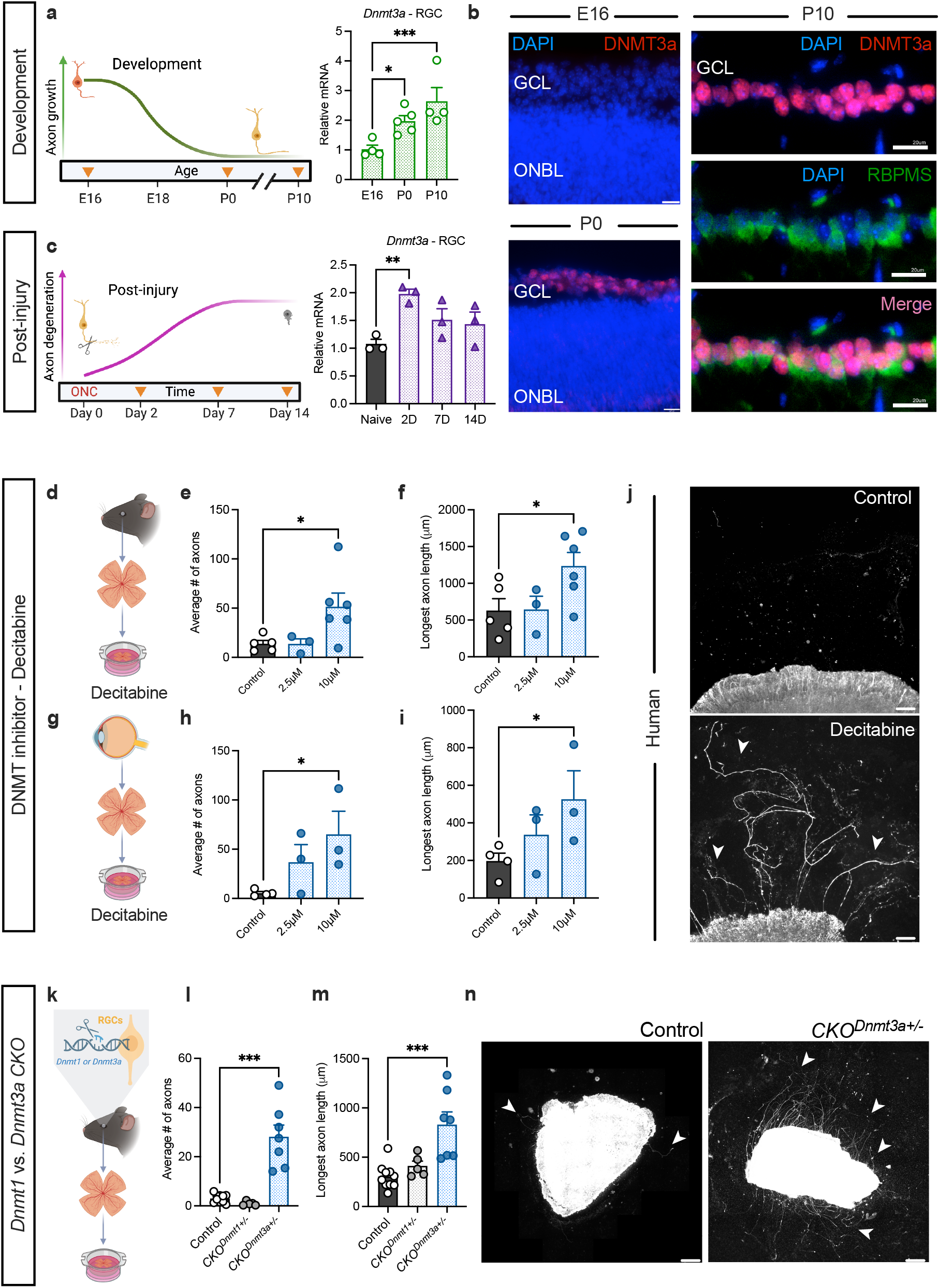
*Dnmt3a* deficiency enabled robust axonal outgrowth in retinal explants of mice and human. **a**, Schematic of RGC axon growth timeline during development (left) and qPCR quantification of *Dnmt3a* expression in purified RGCs (right) (n = 4 – 5 mice/group); note the inverse correlation of *Dnmt3a* expression and RGC axon growth capacity from E16 to P10. **b**, Representative images of retinal sections from mice aged E16, P0, and P10 that were immunolabeled for DNMT3a (red) and an RGC marker RBPMS (green) and counter-stained with DAPI (blue). DNMT3a signal was first detected in the ganglion cell layer (GCL) at P0 and became intense and colocalized with RBPMS in P10 retinas. Scale bar: 20 µm. **c**, Schematic of axon degeneration timeline (left) and qPCR quantification of *Dnmt3a* expression in purified RGCs (right) from naïve and post-ONC mice (n = 3 mice/group). **d-i**, Schematics of experiments presented in **e** and **f** (**d**) or **h-j** (**g**), and quantification of axon number (**e**, **h**) and longest axon length (**f**, **i**) in retinal explant cultures derived from adult mice (**e**, **f**) and post-mortem human eyes (**h**, **i**) that were treated with various doses of a pan DNMT inhibitor decitabine (n = 3 – 6 mice/group). **j**. Representative images of cultured human retinal explants treated with vehicle (control) or decitabine (bottom) and immunolabeled for βIII-tubulin to reveal growing axons. Scale bar: 100 µm. **k-m**, Schematic of experiments presented in **l-n** (**k**) and quantification (**l**, **m**) of axon number (**l**) and longest axon length (**m**) in cultured retinal explants taken from mutant and littermate control (*Cre* and *fl/+*) mice (n ≥ 5 mice/group). **n**, Representative images of cultured retinal explants taken from littermate control (left) and *CKO^Dnmt3a^* (right) mice that were immunolabeled for β-III tubulin. Scale bar: 400 µm. ONBL, outer neuroblastic layer. White arrows denote neurites (\**P* < 0.05, \*\**P* < 0.01, \*\*\**P* < 0.001, one-way ANOVA; mean ± s.e.m.)

To ascertain whether DNMTs affect RGC axonal regrowth, we assessed the effects of decitabine, a commercially available pan DNMT inhibitor^10^ *in vitro.* The antagonistic effect of decitabine on DNMTs in post-mitotic neurons in the retina and brain of adult mice has been well documented^11-13^. We thus applied decitabine to retinal explant cultures, where the growth of axons mirrors the regenerative capacity of axotomized optic nerve fibers *in vivo*^5,6^. Consistent with the lack of nerve regeneration, retinal explants of adult (> 2 months old) wildtype mice and post-mortem human eyes showed minimal axonal outgrowth as revealed by β-III tubulin immunolabeling. In contrast, both mouse and human retinal explants treated with decitabine exhibited significant increases in the number and length of axonal growth (**Fig. 1d-j**).

We next adopted a genetic approach to determine which DNMT(s) regulate axon growth capacity in postnatal RGCs. Two DNMT families play different roles in methylation: DNMT1 maintains the methylation patterns, whereas the DNMT3 family (especially DNMT3a and DNMT3b) establishes initial methylation patterns^14^. We generated conditional knockout mice carrying RGC-specific allele with abolished catalytic activity of *Dnmt1* (*CKO^Dnmt1^*)^15^ or *Dnmt3a* (*CKO^Dnmt3a^*)^16^. *Vglut2-Cre* transgenic mice were previously shown to drive expression of *Cre* recombinase in virtually all RGCs and sporadically cone photoreceptors, but not other retinal cell types^17,18^. We verified this by crossing *Vglut2-Cre* transgenic mice with the *Ai9* mouse line, which expresses the fluorescent reporter tdTomato from the Rosa26 locus in a *Cre*-dependent manner (*R26-tdTomato*). Approximate 90% of RBPMS+ RGCs and a small number of cones^19^, but not other retinal cells, were positive for tdTomato (**Extended Data** Fig. 1d, e).

By generating *CKO^Dnmt1^* and *CKO^Dnmt3a^*, we noted that adult homozygous *CKO^Dnmt3a-/-^* mice developed obesity and malocclusion, presumably because *vGlut2* drives *Cre* expression in many CNS neurons. The mice also suffered a high mortality rate from anesthetics during surgical procedures. In contrast, heterozygous *CKO^Dnmt3a+/-^* mice grew and bred normally without apparent health concerns or retinal structural changes. This prompted us to focus on *heterozygous CKO* mice. We isolated RGCs using magnetic bead-conjugated Thy-1 antibody and detected significant downregulation of *Dnmt3a* mRNA in heterozygous *CKO^Dnmt3a+/-^*RGCs compared to those from the littermate controls (**Extended Data** Fig. 1f). We noted that in homozygous *CKO^Dnmt3a-/-^* RGCs, a low level of *Dnmt3a* mRNA was detected, likely because Thy-1 is not exclusively expressed by RGCs in the rodent retinas^20,21^. We confirmed that *Dnmt1* and *Dnmt3a* mRNA levels were downregulated selectively in the RGCs, but not the entire retinas of *CKO^Dnmt1+/-^* and *CKO^Dnmt3a+/-^* mice, respectively (note that RGCs comprise ∼1% of all retinal cells; **Extended Data** Fig. 1f-i). We studied axonal growth capacity using retinal explant cultures derived from adult *CKO^Dnmt1+/-^* and *CKO^Dnmt3a+/-^*mice, and littermate controls that carried *Cre* without *floxed* alleles or *floxed* alleles without *Cre* (**Fig. 1k-n**). While the retinal explants of control mice showed minimal axonal growth, those from adult *CKO^Dnmt3a^*mice exhibited an over 20-fold increase in axon number (**Fig. 1l**) and at least a 2-fold increase in average axon length (**Fig. 1m**) compared to control mice. The retinal explants of C*KO^Dnmt3a+/-^*and C*KO^Dnmt3a-/-^* mice revealed similarly increased number and rate of axon regeneration (data not shown), suggesting that the absence of one *Dnmt3a* allele is sufficient to unleash the barrier to axon regeneration. In contrast, retinal explants of *CKO^Dnmt1+/-^* mice showed no significant improvement in axonal growth compared to controls (**Fig. 1l and m, Extended Data** Fig. 1j). We further verified that *Dnmt3a* deficiency in *CKO^Dnmt3a+/-^* RGCs did not have significant impact on the expression of *Dnmt1* (**Extended Data** Fig. 1k) or *Dnmt3b* (**Extended Data** Fig. 1l). In agreement with the downregulation of *Dnmt3a*, reduced DNA methylation was detected in *CKO^Dnmt3a+/-^* RGCs, but not other retinal cell types compared to controls (**Extended Data** Fig. 1m, n). We concluded that *Dnmt3a*, rather than *Dnmt1*, negatively regulates the intrinsic program for RGC axon growth during maturation.

### Optic nerve reinnervation and restoration of vision by *Dnmt3a* deficiency

Next, we tested *in vivo* if RGC-specific *Dnmt3a* deficiency restores axons’ regenerative capacity in adult RGCs using the optic nerve crush (ONC) injury model. Adult *CKO^Dnmt3a+/-^*mice, as well as their littermate *Cre* or *floxed* control mice, were subjected to ONC. Axonal regrowth was assessed by labeling RGC axons with cholera toxin B subunit (CTB) two days before mice were sacrificed^9,22^. In control mice, CTB-positive axons were observed only proximal to the crush site (**Fig. 2a**) at all time points examined after ONC, confirming the failure of nerve regeneration. In contrast, numerous RGC axons regenerated across the lesion site for long distances in *CKO^Dnmt3a+/-^* mice, with many growing more than 3 mm distal to the lesion by 14 days post ONC (**Fig. 2a**). Notably, the number of regenerated axons was so great that in most cases, it was impossible to count the number of regenerating axons in optic nerve sections. We thus quantified axon regeneration by measuring CTB-labeling immunofluorescent intensity at various distances posterior to the crush site. At all distances, we detected at least 3-fold increase in the intensity of CTB-labeling in *CKO^Dnmt3a+/-^* mice compared to controls; the difference was > 8-fold up to 750 µm distal to the lesion (**Fig. 2b**). Moreover, RGCs of *CKO^Dnmt3a+/-^* mice displayed a significant increase in survival as evidenced by RBPMS-immunolabeling (50% *vs* 12% at 14 days post-ONC. **Fig. 2c, d**). No signs of RGC proliferation were detected as assessed by EdU incorporation assays (data not shown), suggesting that the increased RGC number was not due to the birth of new neurons or RGCs.

**Fig. 2.**
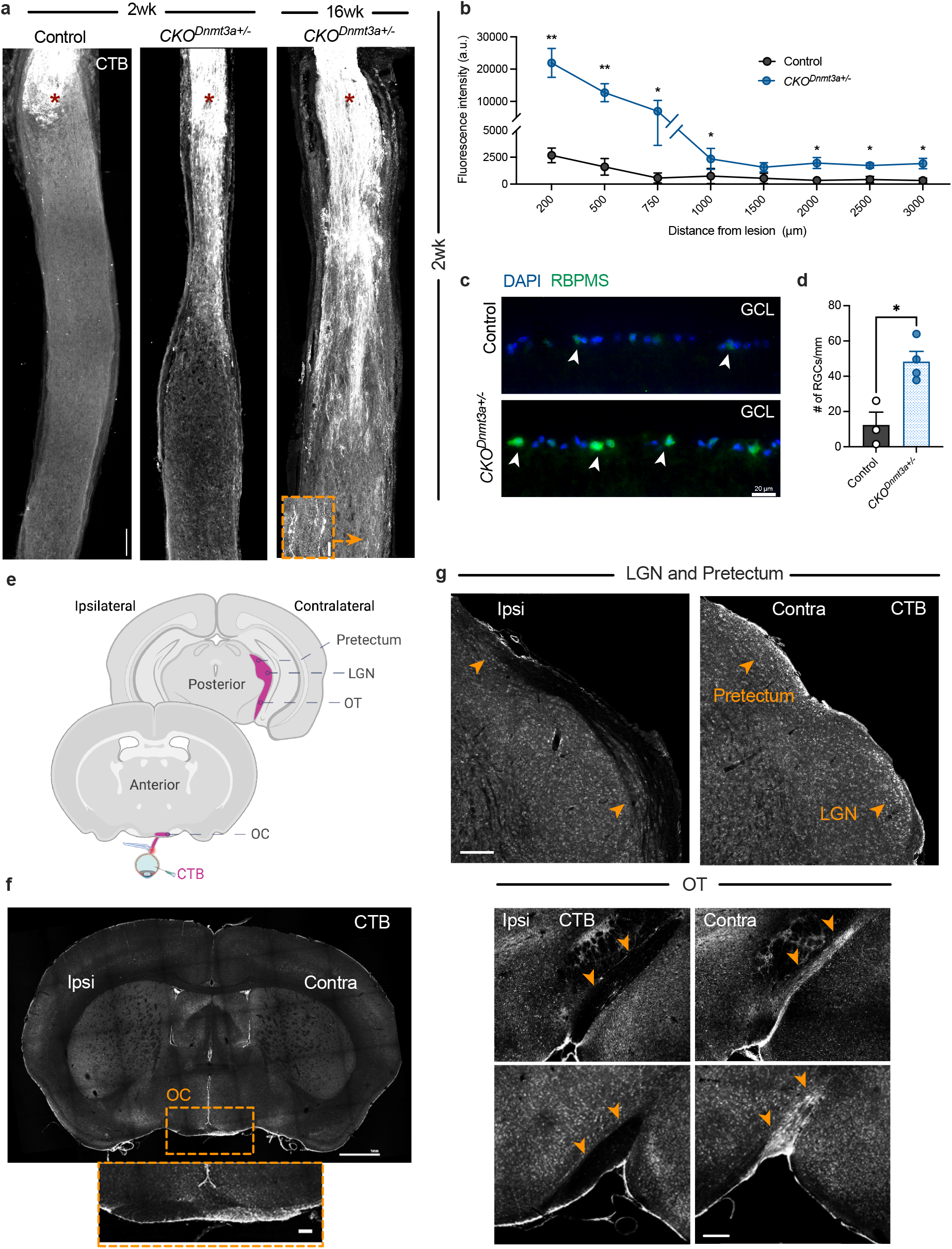
Full-length optic nerve regeneration and target reinnervation after ONC in *CKO^Dnmt3+/-^*mice. **a**, Representative images of longitudinal optic nerve sections showing CTB labeled axons in control and *CKO^Dnmt3a+/-^* mice at 2- or 16-week post-ONC. Each image is an overlay of 3 consecutive nerve sections. Asterisk indicates the crush site. Scale bar: 100 µm; inset: 20 µm. **b**, Quantification of axon regeneration by CTB-labeled fluorescent intensity at various distances distal to the crush site at 2 weeks post-ONC (n = 5 mice/group). **c**, Representative images of retinal sections taken from control and *CKO^Dnmt3a+/-^* mice at 2 weeks post-ONC that were immunolabeled with anti-RBPMS antibody (green) and counter-stained with DAPI (blue). GCL, ganglion cell layer; scale bar: 20 µm. **d**, Quantification of RGC survival by counting RBPMS+ cells in retinal sections at 2 weeks post-ONC. White arrows denote RBPMS+ cells (n = 3 – 4 mice/group). **e**, Schematic of the path of regenerating axons highlighted in purple color, showing their entrance primarily into the contralateral optic chiasm (OC) and optic tract (OT) to innervate the lateral geniculate nucleus (LGN) and Pretectal area. **f-g**, Representative images of brain sections from *CKO^Dnmt3a+/-^* mice at 16 weeks post-crush showing CTB-labeled axons innervating the contralateral (Contra) OC (**f,** scale bar: 1 mm; inset: 100 µm), OT (**g,** scale bar: 200 µm), LGN and pretectum (**g**, scale bar: 200 µm), but not seen in the ipsilateral side (Ipsi). Arrows denote the path of OT, LGN and pretectal areas in Contra and Ipsi brain sections. Insets are circled by orange dash lines. \**P* < 0.05, \*\**P* < 0.01; for **b**, multiple unpaired t-tests; for **d**, unpaired t-test; mean ± s.e.m.

The regenerating axons continued to grow along the optic nerve over time. As early as 4 weeks following the injury, many regenerating axons grew past the optic chiasm, where a major barrier to RGC axon regeneration and target reinnervation was previously reported^23,24^. Greater fluorescent intensity of labeled axons entered the brain when examined at 8- and 16-weeks post-injury, suggesting continual growth of axons. By 16 weeks post-ONC, a large number of regenerating axons crossed the optic chiasm of the contralateral side (**Fig. 2e, f**). In contrast, no axons were evident in the optic chiasm (**Extended Data** Fig. 2a), optic tract, and brain targets (data not shown) of control mice. Unprecedented numbers of regenerating axons were seen in the optic tract of *CKO^Dnmt3a+/-^* mice, primarily extending along the side contralateral to the injury and reinnervating the dorsal and ventral lateral geniculate nuclei (LGN) as well as the pretectum (**Fig. 2g** and **Extended Data** Fig. 2b). These findings show that *Dnmt3a* deficiency in RGCs empowers long-distance and robust optic nerve regeneration and reinnervation into the central visual targets of the optic nerve in adult mice.

We asked if optic nerve reinnervation into the brain targets of adult *CKO^Dnmt3a+/-^* mice led to the restoration of RGC function and vision after ONC (**Fig. 3a**). We assessed mouse visual function by measuring the optomotor response (OMR), which uses the head tracking behavior of mice to determine their spatial vision (e.g., visual acuity)^25^. Normal mice without the injury typically exhibited a visual acuity of 0.4 – 0.5 cycle/degree as measured by OMR. When examined at 2 weeks post ONC, all animals exhibited a complete loss of head tracking behavior in OMR assays, indicative of blindness following nerve injury (**Fig. 3b**). OMR remained absent in control mice throughout the study period (up to 16 weeks post-lesion). In contrast, ∼40% of the *CKO^Dnmt3a+/-^* mice started to show a recovery of OMR by 4 weeks post-injury, a time corresponding to the entry of RGC axons into the brain and reinnervating the central visual targets as documented above. The percentage of *CKO^Dnmt3a+/-^* mice which had regained OMR increased with time and reached ∼70% by 16 weeks post-ONC. Quantification of OMR-based visual acuity reached 0.26 cycle/degree, a recovery of more than 50% of the normal value of visual acuity in mice (**Fig. 3b**).

**Fig. 3.**
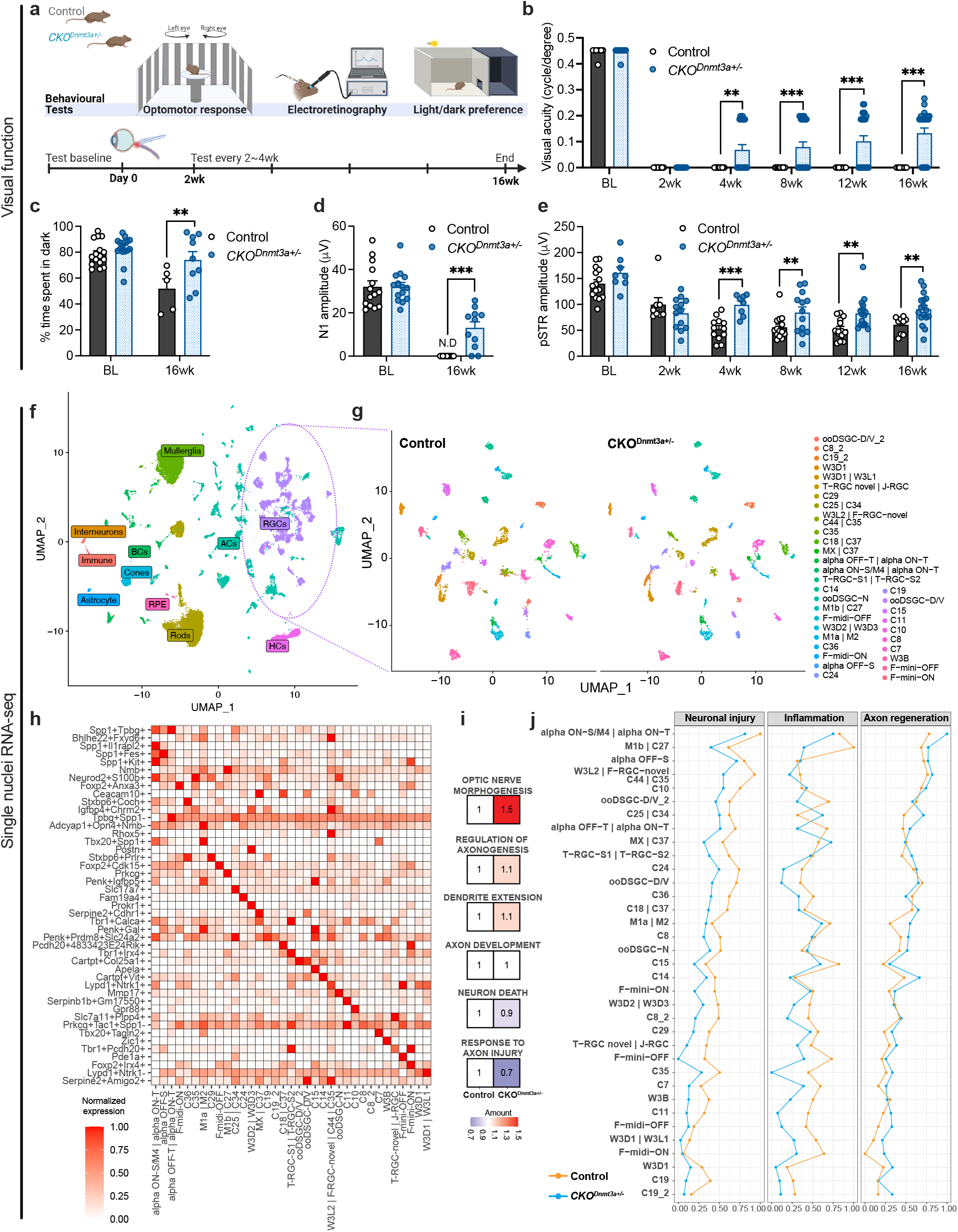
Visual function recovery following ONC in *Dnmt3a* deficient mice. **a**, Schematic of the experiments presented in **b**-**e**. **b**, Quantification of visual acuity assessed by OMR in *CKO^Dnmt3a+/-^* and littermate control mice before (baseline, BL) and at 2 – 16 weeks (wk) after ONC (n ≥ 8 mice/group). **c**, Light perception assessed by the light/dark preference tests in *CKO^Dnmt3a+/-^* and littermate control mice before (BL) and at 16 weeks post-ONC; the assay measured the percentage of time spent in the dark chamber (n ≥ 5 mice/group). **d**, VEP N1 wave amplitudes in *CKO^Dnmt3a+/-^*and littermate control mice assessed before (BL) and at 16 weeks post-ONC (n ≥ 11 mice/group). **e**, RGC function measured by pSTR amplitudes in *CKO^Dnmt3a+/-^* and littermate control mice before (BL) and at 2 – 16 weeks after ONC (n ≥ 8 mice/group) (\**P* < 0.05, \*\**P* < 0.01, \*\*\**P* < 0.001, multiple unpaired t-tests; mean ± s.e.m.). **f**, **g**, Dimplots showing detection of retinal cell (**f**) and RGC types (**g**) in the *CKO^Dnmt3a+/-^* and littermate control retinas (n = 6 retinas from 6 mice/group) at 2 days after ONC by single nuclei RNA-seq (snRNA-seq). **h**, Heatmap showing markers unique for each RGC cluster. X-axis indicates the RGC types (cluster annotation), and Y-axis displays the markers used for RGC type identification. **i**, Single-sample gene set enrichment analysis (ssGSEA) of DEGs from snRNA-seq datasets indicating increased strengths (normalized to controls) in optic nerve morphogenesis, axon genesis, and dendritic extension pathways and decreased strengths in neuronal death and response to axon injury pathways in *CKO^Dnmt3a+/-^* RGCs compared to controls. **j**, Combined GSEA scores in pathways related to neural injury, inflammation, and nerve regeneration among *CKO^Dnmt3a+/-^* (blue dotted line) and control (orange dotted line) RGC types. Normalization of the normalized enrichment scores (NES) was carried out using formulas 4 – 6 described in the Methods. The data revealed across-the-board downregulation of the neuronal injury and inflammation pathways but upregulation of nerve regeneration gene network strength in *CKO^Dnmt3a+/-^* RGCs compared to controls. N.D = not detected.

We performed additional assays to evaluate the RGC and visual functions. First, in a dark/light preference assay, normal mice spent over 80% of their time in a dark environment. Whereas wildtype mice subjected to ONC showed no preference for a dark environment when examined up to 16 weeks post-injury. In contrast, this response in *CKO^Dnmt3a+/-^* mice with ONC recovered to the same level as observed in uninjured mice by 16 weeks post injury (**Fig. 3c**), clear indication of a restoration of light perception. Second, we measured visually evoked potentials (VEPs) in the cortex. Control mice showed an absence of or flat VEP N1 wave (with an infinite delay of N1 latency; **Fig. 3d** and **Extended Data** Fig. 2c, d) up to 16 weeks after nerve crush. Whereas, in parallel with the OMR recovery, ∼60% of *CKO^Dnmt3a+/-^*mice regained VEP N1 responses at an amplitude to half of its baseline value by 16 weeks post-injury (**Fig. 3d; Extended Data** Fig. 2d). Of note, there was a significant delay in N1 latency in *CKO^Dnmt3a+/-^* mice after injury compared to the normal uninjured group (**Extended Data** Fig. 2c, d), potentially due to immature synaptic reconnection or incomplete myelination of the regenerated axons that should impede the speed of electrical signal transmission from the eye to the brain^26,27^. Moreover, significant increases in the amplitude of positive scotopic threshold response (pSTR)^28^, a readout of the electrical activity of RGCs, was detected in *CKO^Dnmt3a+/-^* mice during the period of 4 – 16 weeks post-injury when compared to control mice (**Fig. 3e; Extended Data** Fig. 2e). Together, these results demonstrated reversal of RGC function and vision loss following axon regeneration in a crush-injured optic nerve in adult mice via a single gene manipulation.

### Wide-scale transcriptome shifts by various RGC types towards an injury resilient and regenerative status

Because mouse retina contains 45 RGC types that differ dramatically in their survival and regenerative ability following ONC^18^, it is important to understand if *Dnmt3a* deficiency promotes axon regeneration of specific RGC types. To address the question, we asked if *Dnmt3a* deficiency shifts the RGC landscape or alters the injury responses (pathophysiological condition) in selective RGC types by profiling the RGC transcriptome using single nuclei RNA sequencing (snRNA-seq). Retinas were collected at 2 days post-injury, when few if any RGCs have died but responses to injury are already apparent^5,29^. RGC nuclei were enriched by fluorescence-activated cell sorting (FACS) with a NeuN antibody and profiled by droplet-based snRNA-seq using the 10X platform^30^. We first examined if the proportions of RGC types was affected by *Dnmt3a* deficiency. In the retinas of both *CKO^Dnmt3a+/-^* and control mice, we observed 35 RGC clusters, within which 42 of the 45 reported RGC types were identified based on the RGC atlas markers^18,29^ (**Fig. 3f-h** and **Extended data Fig. 3a-c**). More than half of the clusters (24/35) were 1:1 matches with the reported atlas RGC types, while the other 11 each contained 2-4 RGC types (**Fig. 3h**). The mixture of RGC types within clusters is likely due to the smaller number of RGCs profiled here compared to the earlier report. We found virtually no differences in the distribution of RGC clusters between *CKO^Dnmt3a+/-^* and control mice (**Fig. 3g**). Over 90% of RGC clusters showed similar frequencies in *CKO^Dnmt3a+/-^*and control mice (**Fig. 3f-h** and **Extended Data** Fig. 3a-c). Notably, the frequencies of three αRGC subsets (αON-S/M4|αON-T, αOFF-S and αOFF-T|αON-T), which are the rarest and the most vulnerable and regenerative RGC types^31,32^, were slightly increased in *CKO^Dnmt3a+/-^* mice compared to controls. They collectively represented ∼6% of the total RGC population in *CKO^Dnmt3a+/-^* mice compared to <3% in control mice (**Extended Data** Fig. 3c). Collectively, DNMT3a deficiency does not alter the distribution of retinal cell and RGC profiles or drastically changes the frequencies of the vast majority of RGC types except it somewhat increases the occurrence of rare αRGC subsets.

With this assurance, we assessed differences in gene expression changes after ONC between control and *CKO^Dnmt3a+/-^* RGCs. In line with the robust optic nerve regeneration, we observed upregulation of the nerve growth-related pathways as classified by Gene Ontology (GO) terms, including optic nerve morphogenesis, axonogenesis and dendrite extension (**Fig. 3i**). In contrast, genes associated with neuron death, axon injury response, and neuroinflammation signals classified by GO terms, were downregulated compared to controls, indicating enhanced resilience to injury. By combining the normalized enrichment scores (NES) of injury response- and growth-regulated gene categories, respectively, in each individual RGC clusters, we noted a consistent pattern of gene expression changes across a broad spectrum of RGC types (**Fig. 3j** and **Extended Data** Fig. 3d, 4a, b, 5a**, HTML File**), indicating cell type-independent transformation into growth-promoting and injury-resilient states. Surprisingly, comprehensive downregulation of *Pten* expression, a well-established endogenous inhibitor of axon regeneration which deletion promotes selective axon regeneration of αRGCs^33,34^, was not observed in the majority of *CKO^Dnmt3a+/-^* RGC types, except in four clusters (**Extended Data** Fig. 5b). Moreover, only two *CKO^Dnmt3a+/-^*RGC clusters displayed significantly decreased expression of *Socs3*, another well-known axon regeneration inhibitor^29,34^, when compared to controls. These findings suggest that *CKO^Dnmt3a+/-^* mice switch on a *Pten*- or *SOCS3*-independent axon growth mechanism. Contrary to promoting nerve regeneration in selective RGC types, *Dnmt3a* deficiency triggers a broad-spectrum RGC-transcriptome shift toward a gene profile associated with axon regeneration in nearly all RGC types, including those previously shown to be susceptible to injury (e.g., C35, C25/C34) and those relatively resilient to injury (e.g., αRGCs, M1-RGCs)^18^. Thus, *Dnmt3a* deficiency results in a wide-scale change of transcriptome profiles in various RGC types toward injury-resilient, less inflammatory, and pro-regenerative states after ONC.

### Multifaceted induction of axon regeneration machinery in *Dnmt3a* deficient RGCs

The inability of RGC axons to regenerate is believed to stem from a complex interplay of intrinsic factors and environmental cues, with CNS glial cells releasing axon growth-inhibitory signals and posing a formidable barrier to axon regeneration^1^. To gain deeper insights into the signaling mechanisms through which RGC-specific *Dnmt3a* deficiency disinhibits the development of glial barrier to axonal regrowth, we employed CellChat analysis, a bioinformatic algorithm designed to predict and decipher intercellular cues exchanged by various cell types^35^. Given that DNMT3a dysfunction in *CKO^Dnmt3a+/-^*mice is RGC-specific, we focused on outgoing signals from RGCs to other cell types identified in our snRNA-seq (**Fig. 3f**). As shown above, CellChat analysis consistently predicted significant downregulation of RGC-derived inflammatory signals in *CKO^Dnmt3a+/-^*mice compared to controls, primarily associated with three immune activator (**Fig. 4a** and **Extended Data** Fig. 6a, b): colony stimulating factor (CSF)^28^, granulin (GRN)^36^, and major histocompatibility complex class 1 (MHC-1)^37^. Notably, the chord diagram showed that in control mice, CSF originating from the RGCs signaled to multiple other retinal cell classes, especially immune cells, whereas this RGC signal was drastically diminished in the *CKO^Dnmt3a+/-^*mice (**Fig. 4b**). *Dnmt3a* deficiency also led to downregulation of axon growth-inhibitory signals, such as Semaphorin 4 (SEMA4)^38^, and upregulation of synapse and axon growth-promoting signals, angiopoietin (ANGPT)^39,40^ and secreted phosphoprotein 1/osteopontin (SPP1)^41^ (**Fig. 4a**). Therefore, RGC-specific *Dnmt3a* deficiency not only induces an intrinsic pro-regenerative program within RGCs but also enables a permissive environment by sending decreased inflammatory signals and increased growth-promoting signals.

**Fig. 4.**
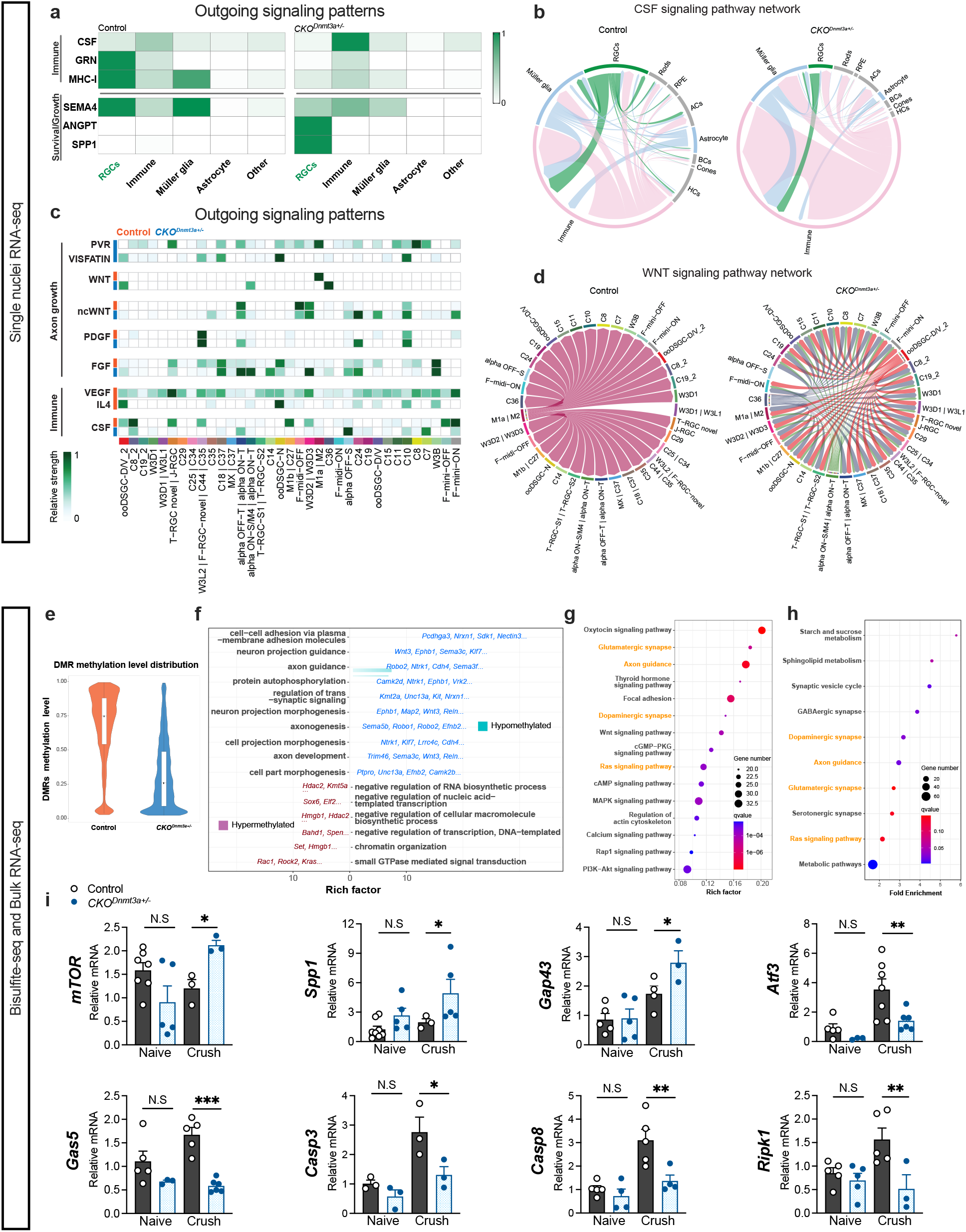
Activation of diverse axon regeneration pathways by *Dnmt3a* deficiency unveiled by multi-omics sequencing data. **a**, Heatmap of cellular outgoing signals in the retina of *CKO^Dnmt3a+/-^* and control mice, predicted by the CellChat output analyses and selected from those showing drastic changes in *CKO^Dnmt3a+/-^* and control RGCs: note the downregulation of CSF, GRN, MHC-I and SEMA4 outgoing signals and upregulation of ANGPT and SPP1 outgoing signals in *CKO^Dnmt3a+/-^* RGCs. X-axis denotes cell populations: RGC, immune, Müller glia, amacrine, and others (including horizontal cells, bipolar neurons, astrocytes, retinal pigment epithelial cells, rods, and cones). Y-axis denotes signaling pathways (receptor-ligand pairs). **b**, Chord diagrams representing the intercellular CSF signaling among retinal cell types. The outgoing signal of RGCs was highlighted in green, while the outgoing signals from all other cell types were colored in gray or light pink to simplify the presentation. Arrows indicate the directions of the signals, and the arrow’s width represents the relative signal strength. **c**, Heatmaps of outgoing signals sent among RGC types in control and *CKO^Dnmt3a+/-^*mice. Note that *CKO^Dnmt3a+/-^* RGCs displayed exclusive transmission of PVR and VISFATIN signals that were absent in control RGCs, while VEGF and IL-4 outgoing signals were observed solely among control RGC types. **d**, Chord diagrams showing intercellular WNT signaling among RGC types in control and *CKO^Dnmt3a+/-^* mice. **e**, Reduced DNA methylation at the differentially methylated regions (DMR) in *CKO^Dnmt3a+/-^* RGCs compared to controls as assessed by whole-genome bisulfite-seq (n = 2 mice/group). **f**, GO analysis of hyper- and hypo-methylated pathways and related genes. **g**, KEGG pathway enrichment of DMR-related gene pathways, selected from TOP-20. **h**, KEGG pathway analysis based on the upregulated DEGs from bulk RNA-seq transcriptome profiling of *CKO^Dnmt3a+/-^* and control RGCs (n = 3 mice/group). Note that the DMR-pathways identified in **g** and the upregulated signaling events in **h** are highly correlated – the common pathways found in both are highlighted in orange. **i**, qPCR quantification of selective genes reported to mediate axon regeneration, neural injury and inflammation in *CKO^Dnmt3a+/-^* and control RGCs isolated from naïve mice or mice at 2 days after ONC (n ≥ 3 mice/group; \**P* < 0.05, \*\**P* < 0.01, \*\*\**P* < 0.001, one-way ANOVA; mean ± s.e.m.).

To determine how *Dnmt3a* deficiency alters the communication among RGC types, we also compared CellChat signaling among RGCs in *CKO^Dnmt3a+/-^* and control mice. The results indicated the up-regulation of classical *Wnt*^42^ and non-canonical *Wnt*^43^, PDGF and FGF signaling^44^ by many RGC types in *CKO^Dnmt3a+/-^*mice compared to controls (**Fig. 4c** and **Extended Data** Fig. 7a, b). The chord diagram of the WNT communication indicated that intrinsically photosensitive RGC types M1a and M2 are the sole major source of WNT signal in controls, while in *CKO^Dnmt3a+/-^* mice, multiple RGC types upregulate outgoing WNT signaling to other RGCs (**Fig. 4d**). *CKO^Dnmt3a+/-^*RGCs also exhibited distinctive expression of poliovirus receptor (PVR)^45,46^ and visfatin^47,48^, two growth and survival-related genes that were absent in control RGCs. Conversely, the growth factors with pro-inflammatory properties, such as vascular endothelial growth factor (VEGF)^49^ and interleukin 4^50,51^, were highly expressed only in control RGCs but not in *CKO^Dnmt3a+/-^*RGCs (**Fig. 4c**). These results strengthen the conclusion that *Dnmt3a* deficiency in the RGC contributes to distinctive pro-regenerative intercellular signaling and suppressed inflammatory responses.

### DNA methylation of axon growth gene networks regulated by *Dnmt3a*

DNMT mediates DNA methylation to serve biological functions. To investigate how *Dnmt3a* deficiency altered the landscape of DNA methylation in RGCs, we conducted whole genome bisulfite sequencing (WGBS) of RGCs isolated 2 days post-ONC. Compared to controls, *CKO^Dnmt3a+/-^*RGCs showed lower CG methylation levels specifically in the differentially methylated regions (DMRs) (**Fig. 4e**), which are the genomic regions show different methylation status between individuals. DMRs are highly regarded as possible functional regions involved in gene transcriptional regulation^52^. In contrast, minimal alteration in genome-wide DNA methylation level was detected comparing the two groups (**Extended Data** Fig. 8a, b). WGBS disclosed 4,836 CG DMRs, including 4,004 hypo-DMRs and 832 hyper-DMRs (**Extended data Fig. 8c**). Annotation of the hypomethylated DMR-related genes identified 48 pathways in the biological processes, which reflect the aforementioned transcriptomic shifts toward a pro-regenerative state in the *CKO^Dnmt3a+/-^* RGCs. Fourteen of these hypomethylated pathways are linked to axon growth, including axon guidance, axonogenesis and axon development (e.g., *Wnt3, Klf7, Map2*)^38,53,54^, and 6 are related to synaptic process (e.g., *UNC13a* and *Nrxn1*)^55,56^ (**Fig. 4f**). KEGG pathway analysis of total DMR-related genes again depicted distinctive enrichment in pathways related to nerve growth and synaptic process that include key signaling events such as *Wnt*, *Ras*, and *cAMP*, axon guidance, and dopaminergic and glutamatergic synapse (**Fig. 4g**). The data establish that DNMT3a directly regulates the methylation of gene networks that control axon growth and synaptic process.

To validate that hypomethylation of genes controlling axon growth and synaptic process in *Dnmt3a*-deficient mice results in correspondent gene expression changes, we conducted comprehensive transcriptome profiling by bulk RNA-seq on RGCs isolated two days after ONC. Principal component analysis clearly separated *CKO^Dnmt3a+/-^* RGCs from the controls (**Extended Data** Fig. 8d). We identified 1,459 genes that are differentially expressed (DEGs) (Fold change > 1.5; padj < 0.05) between *CKO^Dnmt3a+/-^* and control RGCs (**Extended Data** Fig. 8e). KEGG analyses revealed alignment between upregulated gene pathways in *CKO^Dnmt3a+/-^* RGCs with those found to be hypomethylated above, including synaptic signaling and axon guidance (**Fig. 4h**). On the other hand, downregulated DEGs in the *CKO^Dnmt3a+/-^*RGCs were primarily involved in cell death and immune regulation, such as apoptosis, TNFα and chemokine/cytokine signaling (**Extended Data** Fig. 9a). GSEA and GO enrichment studies confirmed the results of KEGG analysis, showing downregulation of immune response (e.g., T cell and macrophage migration, chemokine-mediated signaling, and microglial activation) and inflammatory gene pathways (e.g., IFNγ, TNFα, and IL6-JAK-STAT3 signaling)^57^ with concurrent upregulation of key events regulating axon growth (e.g., *Wnt/*β*-catenin*) and synaptic assembling and transport (**Extended Data** Fig. 9b, c). This result was congruent with the prediction of CellChat analysis.

We further verified the gene expression pattern changes detected by bulk RNA-seq in *CKO^Dnmt3a+/-^* and control mice by qPCR. *CKO^Dnmt3a+/-^* RGCs isolated at 2 days post-injury showed consistent significant differences in the expression of the aforementioned gene pathways, including the upregulation of axon growth-related genes (e.g., *mTOR*, *Spp1*, *Gap43*) and downregulation of growth inhibitors (e.g. *Atf3*) as well as cell apoptotic and inflammatory genes (e.g., *Gas5*, *Casp3*, *Casp8*, *Ripk1*)^1,29,57,58^ compared to control RGCs (**Fig. 4i**). Interestingly, these differences were only detected in RGCs isolated post ONC. RGCs taken from uninjured *CKO^Dnmt3a+/-^* mice showed no significant differences in expression of any of these genes compared to RGCs from uninjured control mice (**Fig. 4i**). The data suggest that the loss of DNMT3a driven by *Vglut2-Cre* has little impact on gene expression of normal RGCs but rather results in reprogramming of RGCs’ injury responses toward an injury-resilient and pro-regenerative status post-ONC. Together, these studies demonstrated that *Dnmt3a* deficiency reshaped the injury-induced transcriptomic and intercellular communication landscape of RGCs, contributing to the functional regeneration of the optic nerve.

### Therapeutic inhibition of *Dnmt3a* rescued visual function in adult wildtype mice after ONC

To assess the therapeutic potential of *Dnmt3a* inhibition, we conducted AAV treatment experiments in adult mice after ONC (**Fig. 5a**). To downregulate *Dnmt3a* we injected AAV-shRNA intravitreally to adult wildtype mice or AAV-Cre to mice carrying heterozygous floxed *Dnmt3a* allele (*fl/+*) immediately after ONC. Detection of GFP expression revealed wide-spread AAV-infection in RGCs, and qPCR analysis confirmed the downregulation of *Dnmt3a* mRNA levels in RGCs of eyes injected with AAV-shRNA or AAV-Cre compared to AAV-scrambled RNA or AAV-GFP injected eyes (**Extended data fig. 9d, e**). Tracking of the spatial vision with OMR at multiple timepoints post-injury indicated partial recovery of visual acuity in over 50% of mice received AAV-shRNA treatment starting at 6 weeks post-injury (**Fig. 5a, b**), a two-week delay of recovery than that was seen in *CKO^Dnmt3a+/-^*mice above. This corresponds to the two week-time period that is required for AAV to reach its peak/plateau of gene expression after intravitreal injection (not shown). In contrast, control mice received injection of AAV-scrambled RNA showed no response in the OMR test up to 12 weeks after injury. By 12 weeks post-ONC, AAV-shRNA-treated mice regained light perception and exhibited no difference in light/dark box assays compared to their uninjured baselines (**Fig. 5c**). Moreover, AAV-shRNA-treated mice demonstrated improved pSTR and N1 amplitude of VEP compared to control mice receiving AAV-scrambled RNA injection, which remained blind without any sign of light perception or VEP response (**Fig. 5d-f**). Similar recovery of visual acuity, light perception, pSTR, and VEP improvement were also observed in *fl/+* mice received AAV-Cre compared to AAV-GFP injected mice (**Fig. 5g-m**). Similarly, delays in VEP N1 latency were observed in both AAV-shRNA-treated wildtype mice and *AAV-Cre*-injected *fl/+* mice, suggesting impaired myelination. These results provide compelling evidence that *Dnmt3a* inhibition holds immense therapeutic potential for the treatment of optic nerve diseases or injury, offering a viable avenue for restoring vision *in vivo*.

**Fig. 5.**
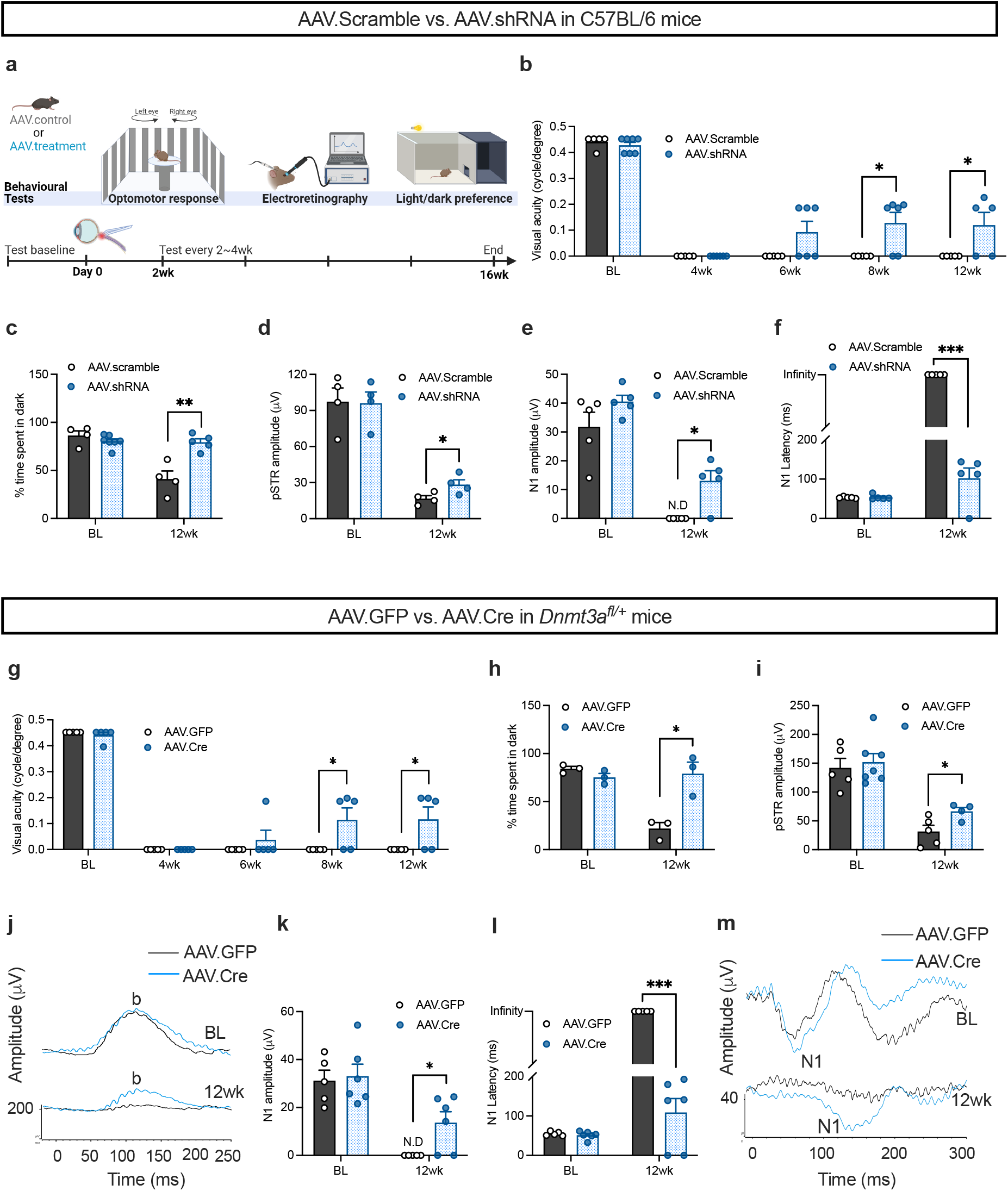
Restoration of vision following ONC in adult mice by therapeutic *Dnmt3a* knockdown. **a**, Schematic of the experiments presented in **b-m**. **b**, Assessment of visual acuity by OMR assay in adult wildtype mice before (BL) and at 4 – 12 weeks after ONC; mice received intravitreal injection immediately after ONC of either control *AAV.scramble* or treatment *AAV.shRNA* that specifically targeted *Dnmt3a*. **c**, Percentage of times spent in the dark chamber by mice in the light/dark preference test. **d**, pSTR amplitudes. **e**, **f**, VEP N1 amplitude (**e**) and latency (**f**). **g**, Assessment of visual acuity by OMR assay in adult *Dnmt3a^fl/+^*mice before and at 4 – 12 weeks after ONC; mice received intravitreal injection immediately after ONC of either control *AAV.GFP* or treatment *AAV.Cre*. **h**, Percentage of time spent in the dark chamber by mice in the light/dark preference test. **i**, pSTR amplitude. **j**, Representative pSTR wave forms of (**i**). **k**, **l**, VEP N1 amplitude (**k**) and latency (**l**). **m**, Representative VEP waveforms of (**k**, **l**). (n ≥ 3 mice/group; \**P* < 0.05, \*\**P* < 0.01, \*\*\**P* < 0.001, multiple unpaired t-tests; mean ± s.e.m.).

## Discussion

Here we report restoration of vision after traumatic injury of the optic nerve via suppression of *Dnmt3a*, which can overcome the inability of adult RGCs to regenerate axons. Inhibition of DNMT3a-dependent DNA methylation reactivates an intrinsic axon growth program that is lost in adults, promotes RGC survival, and enables axonal regeneration through the optic nerve to central targets. As regenerating axons reach the brain, spatial vision is recovered. This finding highlights the dominant role played by neuron-intrinsic determinants in driving functional optic nerve regeneration and alleviating environmental obstacles, likely through signaling from RGCs to other cell classes and to each other. Moreover, *Dnmt3a* deficiency facilitates selective demethylation of gene networks and multifaced activation of axon growth machinery across diverse RGC subtypes without apparent neuroinflammation or compromised neuronal survival. Most importantly, therapeutic suppression of *Dnmt3a*, either by AAV-shRNA delivery or other means, holds the potential for clinical translation. This discovery not only offers an exciting avenue for treating injury to optic nerve diseases and injury, but also paves the way for future research in brain and spinal cord regeneration.

## Supporting information

Extended Figures with Legends

Extended Data HTML

## Acknowledgements

This work was supported by the National Eye Institute Grant R01 EY031696, R21 EY033882 and a Harvard NeuroDiscovery Center Grant to D.F.C., a Core Grant for Vision Research from NIH/NEI to the Schepens Eye Research Institute (P30EY003790), and R21 EY032219 to J.R.S. We thank Haobo Li for plasmid construction and AAV production; Jenna Cho, Alice An, and Irvine Yi for assistance in data quantification; Anton Lennikov, Shuhong Jiang, and Karen Chang for advice and assistance in imaging and visual function studies; Kasim Gunes and Julie Chen for support in culture studies. We thank the Schepens Eye Institute Animal Care Facilities for support in the animal studies, the Gene Transfer Vector Core, and the Morphology Core facility, which is supported by the NIH-National Eye Institute P30 Core Grant (P30EY003790).

## Author contributions

D.F.C and W.L.T. perceived the project, analyzed the data, and wrote and revised the manuscript with input from all co-authors. W.L.T. was involved in all experiments and analyses. K.C. contributed to the optic nerve crush studies and trained W.L.T. in all animals works and analyses. E.K. designed and performed all sequencing data computational analysis. A.M. performed single-nucleus RNA sequencing and W.Y. contributed to single-nucleus data preprocessing. A.A. contributed to the culture studies and assisted with image acquisition. Y.L. and X.W. conducted early small-molecule and genetic screening studies. T.P.G. contributed to qPCR and morphometric analyses. J.R.S and P.B. carefully read and edited the manuscript and supervised single-nucleus RNA sequencing studies.

## Methods

### Mice

*Vglut2-ires-Cre* knock-in (*Vglut2-Cre*) mice were generated in Bradford B Lowell’s laboratory (Harvard University)^1^ and *Dnmt3a* floxed (*Dnmt3a^flox^*) mice were obtained from Guoping Fan’s laboratory (University of California, Los Angeles)^2^. C57BL/6J wildtype (000664) and B6.Cg-Gt(ROSA)26Sortm9(CAG-tdTomato)Hze/J (007909) mice were purchased from the Jackson Laboratory. Both sexes of animals were used in the experiments. The control group for experiments with Cre/lox mice included *Vglut2-Cre*, floxed, and littermate wildtype mice. All animal procedures are approved by the Institutional Animal Care and Use Committees (IACUCs) at the Schepens Eye Research Institute (SERI), Mass Eye and Ear. Animals were bred and housed under 12-h light/dark cycles with food and water ad libitum in the SERI animal facility.

### RGC isolation

As described previously^3^, mouse retinas were dissected in Neurobasal-A medium (Gibco, 10888-022) on ice and dissociated into single-cell suspensions in papain supplemented with deoxyribonuclease I and Ovomucoid protease inhibitor by manual trituration according to the manufacturer’s protocol (Papain Dissociation System, Worthington, Lk003150). The resulting cells were spun down and resuspended in autoMACS Rinsing Solution (Miltenyi Biotec, 130-091-222) followed by incubation with micro-magnetic beads conjugated Thy1.2 antibody (Miltenyi Biotec, 130-121-278) for 20 min at 4°C. Cells were then loaded into the MS Column (Miltenyi Biotec, 130-042-201) held by a magnetic separator (Miltenyi Biotec, 130-042-108) that allowed separation of magnetic beads-labeled cells from the rest by rinsing the column. After 3 washings, the column was removed from the separator and the labeled cells were gently flushed into a collection tube with 1 ml of the rinsing solution. The collected cells were spun down and processed for RNA or DNA analysis.

### Total nucleic acids extraction

Total RNA and DNA were extracted using the Zymo Quick-RNA MicroPrep Kit (Zymo Research, R1051) and Quick-DNA MiniPrep Kit (Zymo Research, D3024) respectively. The quantity and quality of both nucleic acids were measured using a NanoDrop 2000 Spectrophotometer (Thermo Fisher Scientific). RNA was then frozen and sent to Novogene for ultra-low-input bulk RNA sequencing or converted into cDNA with the PrimeScript RT Master Mix (Takara Bio, RR036A) for quantification of mRNA level using qPCR^4^. Primers used for qPCR are listed **in Supplementary Table 1**. DNA was frozen and sent to Novogene for whole genome bisulfite sequencing (WGBS) or processed for assessment of global methylation level.

### Global DNA methylation assay

Genome-wide DNA methylation levels were determined using the 5-mC DNA ELISA Kit (Zymo Research, D5325) per the manufacturer’s instruction. Absorbance was recorded at 405 nm with a microplate reader (Agilent, BioTek Synergy H1 Multimode Reader). The percentage of methylated cytosines (% 5-mC) in the total DNA content was determined by calculating the levels of 5-mC using a standard curve generated with the kit controls.

### Human eye globes

The Lions Eye Bank in Florida provided human cadaver eye globes for the experiments. These eye globes were obtained from donors between the ages of 68 and 92, as stated in **Supplementary Table 2**. All experiments involving human tissue were conducted according to the principles of the Declaration of Helsinki.

### Retinal explant culture

Retinas dissected out from post-mortem human eyes were cut at both central and peripheral areas with a biopsy punch (Acuderm Inc, P2525) in ice-cold Neurobasal-A medium, producing small explants in 2.5 mm diameter. Retinas from adult mice were cut into 4-6 equally sized pieces. With the RGC layer facing down, retinal explants were placed onto tissue culture inserts (6-well plate format, Greiner Bio-One, 657641) that were pre-coated with Matrigel matrix (Corning, 354230). Inserts were then placed into culture plate wells containing Neurobasal-A medium supplemented with 25 µM L-Glutamic acid (Sigma-Aldrich, G8415), 2 mM Glutamax (Gibco, 35050061), 1× penicillin/streptomycin (Thermo Scientific, 15140122), 1× B-27a (Gibco, 17504-044), 5 µg/ml insulin (Sigma-Aldrich, I9278), 50 ng/ml BDNF (PeproTech, 450-02), 50 ng/ml CNTF (PeproTech, 450-13), and 1× forskolin (Sigma-Aldrich, F6886). Explants were cultured at 37°C with 5% CO_2_ for 7 days and then processed for immunohistochemical analysis.

### Quantification of neurite outgrowth in explant cultures

After 7 days of incubation, retinal explants on inserts were fixed with 4% PFA for 2 h and blocked with Mojito buffer (10% NGS, 3% NDS, 1% BSA, 0.5% Tween-20, 0.5% Triton X-100 and 0.1% sodium citrate buffer in PBS) for 2 h at room temperature. To stain for neurites, explants were incubated with mouse anti-β Tubulin III (TUJ1) antibody (1:500, Novus Biologicals, NB600-1018) overnight at 4°C followed by Alexa Fluor 594-conjugated secondary antibody (1:500, Jackson ImmunoResearch) for 2 h at room temperature. All antibodies were diluted in a staining buffer consisting of 0.5% Triton X-100, 0.1% Tween-20 and 5% BSA in PBS. Between changes of solution, explants were washed 3 times, 10 min each with PBS. After mounting onto Superfrost Plus Slides (VWR, 48311-703) with Fluoromount-G (SouthernBiotech, 0100-20), explants were imaged with a Leica fluorescence microscope (Leica Microsystems, DMi8). The neurite number and length of each retinal explants were quantified using the built-in software LAS X of the microscope (Leica Microsystems, Leica Application Suite X) by individuals blinded to the experimental groups.

### Optic nerve crush injury

Mice were anesthetized by inhalation of 3-4% isoflurane for induction and 1-3% isoflurane for maintenance. Detailed surgical procedure of unilateral optic nerve crush (ONC) has been described previously^5,6^. In brief, a small incision was made in the temporal conjunctiva of the mouse eye, and the optic nerve was exposed and crushed with a Dumont #5 jeweler’s forceps (FST) for about 5 sec, approximately 0.5 mm behind the eyeball. Proparacaine and antibiotic ointment were applied to the ocular surface before and after the crush, respectively. Buprenorphine SR or Ethiqa X were given subcutaneously as a postoperative analgesic.

### Immunohistochemistry of cryosections

Mouse eye and brain were dissected out after transcardiac perfusion with 4% PFA in PBS. Tissues were post-fixed in 4% PFA overnight and then cryoprotected in 30% sucrose overnight at 4°C. For cryosections of the eye, cornea and lens were removed from the eye before being embedded in OCT compound (VWR, 25608-930) and frozen. The frozen tissue block was then sliced into 16 μm serial cross sections, collected on Superfrost Plus Slides and stored at -20°C until processed. Frozen sections were blocked with BSA (5%) and Triton X-100 (1%) in TBS for 1 h at room temperature, followed by incubation with the primary antibodies overnight at 4°C and then the proper secondary antibodies (1:500, Alexa Fluor 488, 594, 647 conjugated; Jackson ImmunoResearch) for 2 h at room temperature in the same blocking solution. Between changes of antibody incubation, all sections were washed 3 times, 5 min each. For staining of cholera toxin subunit B (CTB), after primary antibody, sections were incubated with biotin anti-goat (1:250, Vector Laboratories, BA-9500) for 2 h and Alexa Fluor 594-conjugated streptavidin (1:500, Thermo Scientific, S32356) for 1 h at room temperature. Primary antibodies used are goat anti-CTB (1:4000, List Biological Laboratories, 703), guinea pig anti-RBPMS (1:200, PhosphoSolutions, 1830-RBPMS), and mouse anti-DNMT3a (1:100, Novus Biologicals, NB120-13888SS). Staining for anti-DNMT3a required 2N HCl treatment for 30 min at 37°C followed by neutralization with 0.1M Trish-HCl (pH 8.3) for 10 min at room temperature before the blocking.

For cryosections of the brain, OCT-embedded tissue block was cut into 40 μm serial cross sections, collected in cryoprotectant solution and stored at -20°C until processed. The floating brain sections were blocked with Background Buster (Newcomer Supply, NB306-50) for 30 min at room temperature followed by a previously published staining protocol with slight modification. Briefly, after 4-day primary incubation with goat anti-CTB (1:4000) at 4°C, floating sections were stained with biotin anti-goat for 2 h (1:200) and Alexa Fluor 594-conjugated streptavidin (1:1000) for 2 h at room temperature.

Immunostained samples were mounted with Fluoromount-G and imaged using Leica DMi8 fluorescence microscope and Leica SP8 confocal microscope (Leica Microsystems) with 20x and 40x lens, and Olympus FluoView FV3000 confocal laser scanning microscope (Olympus Life Science) with 4x lens. Tile scans of 3 – 4 consecutive retinal and longitudinal optic nerve sections per animal were imaged and used for RGC and axon quantification where data from the scans were averaged for individuals. Projection of Brain Z-scans was carried out using the ImageJ (1.53t) Fiji (2.13.1) with the ‘Average intensity’ projection option. Quantification of regenerated axons was adapted from protocols described previously^7^. In brief, within the microscope’s built-in software LAS X, the fluorescence intensities of CTB-positive axons at different distances post the crush site were measured as the integrated density of the Region of Interest (ROI) subtracting the background. For quantification of RGC density in retinal sections, retinal sections containing the optic nerve head (thus crossing over the central and peripheral retinal regions) were used. The total number of RGCs in each retinal section was recorded and divided by the length of the entire retinal section. The quantifications were performed by individuals blinded to the experimental conditions.

### Optomotor response-based visual acuity test

The optomotor reflex-based spatial frequency threshold test was performed as described previously to measure the visual acuity of mice^4^. A mouse was placed on a pedestal in the center of an area surrounded by four LCD screens (Acer 15-inch) that displayed black and white stripes in a grating pattern, which rotated clockwise or counter-clockwise. Each eye was tested separately based on the direction of the rotating stripes. Two observers watched the mouse’s tracking behavior, and a positive response was determined when the mouse head followed the grating movement. The rotation speed and contrast were kept constant. The width of the bars was varied to test how fine the mouse could see, and the corresponding bar width was calculated in term of cycle/degree where 0.4∼0.5 cycle/degree reflects normal visual acuity in mouse. The responses were measured before and after treatment by individuals blinded to the group or treatment of the mouse. Mice with bleeding or inflammation after surgical procedures were excluded from the analysis. The exclusion criteria were established before the experiment.

### Light/dark preference test

The light/dark preference test is based on the propensity of mice to prefer dark over bright environments^8^. A mouse was placed in a box with a light and a dark chamber connected by a small opening to allow free transitions between these two chambers. Mice that can see light show a preference for the dark chamber while blind mice would have no preference. No eyes were covered during baseline measurement while the contralateral eyelid was sutured to prevent the uninjured eye from seeing when tested after crush. Each mouse was kept in the box for 5 minutes and the time it spent in the dark chamber was calculated as a percentage of the total time in the box. In a preliminary study, we showed that mice with both eyes sutured exhibited no preference for the dark chamber as the blind mice, indication of effective blockade of light perception by suturing the eye as it was reported^9^.

### Electroretinography and visual evoked potential

Electroretinography of positive scotopic threshold response (pSTR) and visual evoked potential (VEP) were measured as described previously^4,10^. For pSTR recording, mice were dark adapted for 6 – 12 h before experiment started; while for VEP recording, mice were briefly adapted to dim red light (5 – 10 min) beforehand. Mice were anaesthetized with ketamine/xylazine (100 mg kg−1 and 20 mg kg−1). A drop of 1% tropicamide ophthalmic solution (Bausch & Lomb Inc., Tampa, FL, USA) was applied to dilate the pupils. Dim red light was on throughout the procedure and mice were kept on a built-in warming platform in the Ganzfield ColorDome (ColorDome LabCradle mouse ERG testing, Diagnosys LLC) to prevent hypothermia. Gold wire electrodes contacting both corneas were used to record pSTR with a reference and a ground needle electrode inserted subcutaneously between the eyes and at the base of the tail, respectively. Lights at intensities 6.57E-5 cd.s/m^2^ and 1.7E-4 cd.s/m^2^ were flashed in the dome to elicit pSTR per intensity. An average of 40 responses were recorded by the ERG system (Espion Electroretinography System, Diagnosys LLC) and the b-wave amplitude was measured from the baseline to the positive peak. For VEP, electrode needles were placed subcutaneously at the snout, between the ears, and at the base of the tail. The contralateral eye was covered with a dark patch when recording the ipsilateral eye and vice versa. VEP was stimulated by 100 flashes at 3.0 cd.s/m^2^ and the averaged response was recorded by the ERG system.

### Single-nuclei RNA sequencing

Single-nuclei RNA sequencing was performed based on a previously described method^11^. Retinas from the ONC eyes were collected from 6 control and 6 *CKO^Dnmt3a+/-^* mice, respectively, at 2 days after injury (1 eye/mouse). To extract nuclei, frozen retinal tissues were homogenized in a Dounce homogenizer in 1 ml NP-40 lysis buffer and pelleted at 500 rcf for 5 min. The nuclei were stained for RBFOX3/NeuN (1:300, Sigma, #FCMAB317PE or #MAB377A5) and PTPRC/CD45 (1:300, BD Pharmingen, clone 30-F11) to enrich RGCs and immune cells, respectively. NeuN+ and CD45+ single nuclei were collected in separate tubes using a flow cytometer, and again pelleted at 500 rcf for 5 min. The nuclei were resuspended in 0.04% non-acetylated BSA/PBS solution, adjusted to a concentration of 1000 nuclei/µL, and loaded into a 10X Chromium Single Cell Chip (10X Genomics, Pleasanton, CA) with a targeted recovery of 8000 nuclei. As a result, we sequenced a total of 17,895 nuclei in the control group (Neun+, 9210; CD45+, 8685) and 16,683 nuclei in the *CKO^Dnmt3a+/-^* group (Neun+, 8235; CD45+, 8448). Single nuclei libraries were generated using the Chromium 3’ V3.1 platform (10X Genomics, Pleasanton, CA) following the manufacturer’s protocol, and sequenced on an Illumina NovaSeq at the Bauer Core Facility at Harvard University.

### Single-nuclei RNA-seq data analysis

Data were processed in Cell ranger^12^ to generate the output folder with barcodes, matrix and genes files. Those files were used to generate Seurat object using CreateSeuratObject() function in Seurat package^13,14^. We filtered data to remove doublets, low quality cells and empty droplets and performed filtering based on features/counts/mitochondrial genes/ribosomal genes following standard Seurat pipeline^15^. After data quality control and filtering, a total of 34,102 nuclei from both experimental groups were used for downstream analysis. Sequentially, data were normalized using the “NormalizeData” method, 2000 highly variable features were selected, data were centered, scaled, and clustered with Louvain algorithm^16^. UMAP was chosen as a non-linear dimensionality reduction approach^17^.

We utilized DimPlot() and FeaturePlot() functions in Seurat as visualization methods for the data annotation. By obtaining the differentially expressed genes list with the function FindAllMarkers(only.pos = TRUE, min.pct = 0.25, log2FC.threshold = 0.25), we performed manual annotation for the dataset clusters (object$seurat_clusters) upon integrating the conditions (Control and *CKO^Dnmt3a+/-^*). The integration was performed using the IntegrateData() function available in Seurat v4. We identified retinal ganglion cells (RGCs), immune cells, Müller glia, amacrine cells, horizontal cells, bipolar neurons, astrocytes, retinal pigment epithelium, rods, and cones, using expression of genes known to be selective markers for each cell class^18^. The immune cells population contained of microglia/macrophages (Klra17+, C1qc+, Cx3cr1+, Tmem119+), T helpers (H2-Ab1+, Cd74+), T cells (Grap2+, Itgb7+). As the total number of immune cells in the populations was small, we labeled them as ‘Immune’ for the downstream analysis. FindAllMarkers() function was used to identify the gene expression dynamics between the conditions on total RGC subset of the dataset.

We reclustered RGCs, normalized them using SCTransform method^19^ and used previously identified type-specific markers for RGC types identification^20^. With the type markers list, we utilized AverageExpression() Seurat function to obtain the gene expression matrix and visualized it using DotPlot() function. Upon identification based on the gene expression, we annotated 35 clusters that contained 42 out of 45 known RGC types^20^. In detail, from those 42 types, we could reach the resolution of 21 types in 24 separate clusters (1:1), with the other 21 types represented in 11 clusters, each of which contained 2 – 4 RGC types (W3D1 | W3L1, M1a | M2, alpha OFF-T | alpha ON-T). The reason we could not reach the resolution of RGC types published by Tran et al^20^ is likely due to the relatively small number of cells available for this analysis.

To visualize RGC type-specific marker gene expression, we generated a matrix of RGC Louvain clusters (object$seurat_clusters) with the expression values for the query gene of interest. We normalized the gene expression ‘n’ from 0 to 1 for every gene k independently per cluster c (formula 1) and generated the values of gene patterns expression p for the gene patterns m (Gene1+Gene2+, Gene1+Gene2+Gene3-, etc.) by summarizing the normalized gene expression values ‘n’ per cluster c (formula 2). Upon receiving the values per pattern showing the gene pattern expression p, those were normalized from 0 to 1 per pattern m independently for every cluster c. The resulting matrix of normalized gene expression patterns ‘N’ was visualized using ggplot2 geom_tile()^21^.

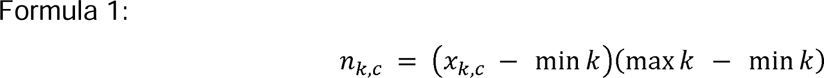

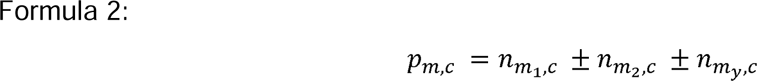

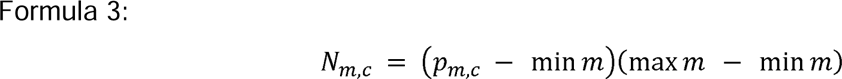

To unravel cell-cell interactions, we utilized CellChat package^22^. For that, we performed two parallel analyses between the conditions: 1) merged RGC clusters vs. every other cluster in the dataset (immune cells, Müller glia, amacrine cells, horizontal cells, bipolar neurons, astrocytes, retinal pigment epithelium, rods, and cones); 2) subset RGC and performed the same type of the analysis for RGC clusters.

The analysis included the CellChat approach to generate total incoming/outgoing signaling and chord diagrams between Control and *CKO^Dnmt3a+/-^*cells with additional settings. Those included type = “truncatedMean”, trim = 0.1, raw.use = FALSE, population.size = TRUE for computeCommunProb() function. Upon generating the data, we merged data of horizontal cells, bipolar neurons, astrocytes, retinal pigment epithelium, rods, and cones and labeled those as ‘rest. For the results, we demonstrated outgoing signaling using pattern = ‘outgoing’ setting for the netAnalysis_signalingRole_heatmap() function and chord diagrams using netVisual_aggregate() function. We exported the computational probability to enable the communications comparisons between the cell types and conditions that were stored in the object as cellchat_object@netP$centr$pathway_name$outdeg and performed 0 to 1 normalization to visualize the data.

To perform GSEA pathway analysis, we utilized the escape package^23^. We loaded C5 pathways from Molecular Signatures Database^24,25^ (‘GOBP_NEURON_DEATH’, ‘GOBP_DENDRITE_EXTENSION’, ‘GOBP_OPTIC_NERVE_MORPHOGENESIS’, ‘GOBP_AXON_DEVELOPMENT’, ‘GOBP_REGULATION_OF_AXONOGENESIS’, ‘GOBP_RESPONSE_TO_AXON_INJURY’, ‘GOBP_INFLAMMATORY_RESPONSE’, ‘GOBP_INFLAMMASOME_COMPLEX_ASSEMBLY’, ‘GOBP_ACUTE_INFLAMMATORY_RESPONSE’, ‘GOBP_CYTOKINE_PRODUCTION_INVOLVED_IN_INFLAMMATORY_RESPONSE’, ‘GOBP_NEUROINFLAMMATORY_RESPONSE’) and performed ssGSEA enrichment^24^. ggplot2 and ggpubr packages^26^ were used for the data visualization.

Then, we generalized the pathways used for GSEA analysis into three programs: neuronal injury, inflammation, and regeneration. The neuronal injury program included ‘GOBP_NEURON_DEATH’ and ‘GOBP_RESPONSE_TO_AXON_INJURY’ pathways. The inflammation program included ‘GOBP_INFLAMMATORY_RESPONSE’, ‘GOBP_INFLAMMASOME_COMPLEX_ASSEMBLY’, ‘GOBP_ACUTE_INFLAMMATORY_RESPONSE’, ‘GOBP_CYTOKINE_PRODUCTION_INVOLVED_IN_INFLAMMATORY_RESPONSE’, and ‘GOBP_NEUROINFLAMMATORY_RESPONSE’ pathways. The regeneration program included ‘GOBP_DENDRITE_EXTENSION’, ‘GOBP_OPTIC_NERVE_MORPHOGENESIS’, ‘GOBP_AXON_DEVELOPMENT’, and ‘GOBP_REGULATION_OF_AXONOGENESIS’ pathways.

To normalize the contribution of the pathways to their corresponding programs from 0 to 1 (due to different average expressions and the amount of the pathways used), we generated the matrix of normalized pathway average expression ‘A’ for every pathway k per cluster c, using the pathway average expression ‘A’ data generated with GSEA analysis approach (formula 4).

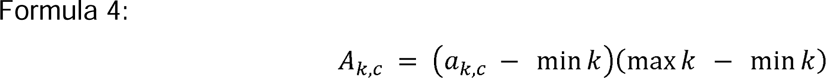

Then, we quantified the pattern average expression ‘m’ for pathway mode y per cluster c as a summation of n pathways average expression A per cluster c with n = 5 for injury, n = 2 for inflammation, and n = 4 for regeneration program (formula 5).

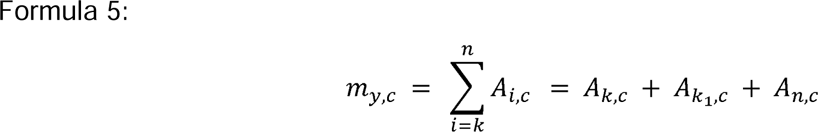

Finally, we quantified the 0 to 1 normalized program average expression ‘M’ for pathway mode y per cluster c as a max/min normalization of program average expression m (formula 6).

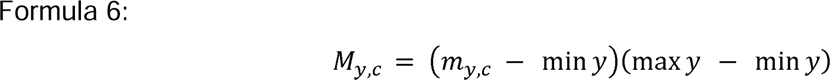

The resulting matrix of RGC subtypes and programs was visualized using geom_point() and geom_line() functions of ggplot2 package with the programs separated using ∼facet_wrap() function. We also performed the 3D visualization for neuronal injury/inflammation/regeneration patterns using plotly package (Extended Data HTML File), and 2D embeddings for injury/regeneration and inflammation/regeneration programs to demonstrate the distribution of RGC subtypes. For 2D and 3D embeddings, the values used are quantified as a difference of normalized program average expression M for pathway mode y per cluster c between *CKO^Dnmt3a+/-^* and Control conditions.

### Whole genome bisulfite-seq analysis

DNA used for bisulfite sequencing were extracted from RGCs isolated from the injured eye at 2 days post-ONC. For each group, RGCs from 2 mice retinas were combined for DNA extraction (n = 2 mice/group, 1 eye/mouse). Bisulfite sequencing and data analysis was performed by Novogene. Basic statistics on the quality of the raw reads was processed by FastQC (fastqc_v0.11.5)^27^. The read sequences produced by the Illumina pipeline in FASTQ format were pre-processed through Trimmomatic (Trimmomatic-0.36) software^28^ using the parameter (SLIDINGWINDOW:4:15; LEADING:3, TRAILING:3; ILLUMINACLIP: adapter.fa: 2: 30: 10; MINLEN:36). The remaining reads that passed all the filtering steps were counted as clean reads and all subsequent analyses were based on this. Finally, we used FastQC to perform basic statistics on the quality of the clean data reads. Bismark software (version 0.16.3)^29^ was used to perform alignments of bisulfite-treated reads to a reference genome (-X 700 --dovetail). The reference genome was first transformed into bisulfite-converted version (C-to-T and G-to-A converted) and then indexed using bowtie2^30^. Sequence reads were also transformed into fully bisulfite-converted versions (C-to-T and G-to-A converted) before they are aligned to similarly converted versions of the genome in a directional manner. Sequence reads that produce a unique best alignment from the two alignment processes (original top and bottom strand) are then compared to the normal genomic sequence and the methylation state of all cytosine positions in the read is inferred. The same reads that aligned to the same regions of genome were regarded as duplicated ones. The sequencing depth and coverage were summarized using deduplicated reads. The results of methylation extractor (bismark_methylation_extractor, - - no_overlap) were transformed into bigWig format for visualization using IGV browser^31^. The sodium bisulfite non-conversion rate was calculated as the percentage of cytosine sequenced at cytosine reference positions in the lambda genome.

To identify the methylation site, we modeled the sum Mc of methylated counts as a binomial (Bin) random variable with methylation rate. To calculate the methylation level of the sequence, we divided the sequence into multiple bins, with bin size is 10kb. The sum of methylated and unmethylated read counts in each window were calculated. Calculated ML was further corrected with the bisulfite non-conversion rate according to previous studies^32^.

Differentially methylated regions (DMRs) were identified using the DSS software^33-35^. The core of DSS is a new dispersion shrinkage method for estimating the dispersion parameter from Gamma-Poisson or Beta-Binomial distributions^36^. DSS possesses three characteristics to detect DMRs. First, spatial correlation. Proper use of the information from neighboring Cytosine sites can help improve estimation of methylation levels at each Cytosine site, and hence improve DMR detection. Second, the read depth of the Cytosine sites provides information on precision that can be exploited to improve statistical tests for DMR detection. Finally, without biological replicate, DSS combines data from nearby Cytosine sites and uses them as ‘pseudo-replicates’ to estimate biological variance at specific locations. According to the distribution of DMRs through the genome, we defined the genes related to DMRs as genes whose gene body region (from TSS to TES) or promoter region (upstream 2kb from the TSS) have an overlap with the DMRs. We quantified the rich factor as the number of DMR-related genes (input genes) divided by the number of pathway genes for each pathway, and the resulting parameters (rich factor, methylation type, and p-value) for the pathways selected from top-20 sorted by p-value (input genes ≥ 20 per pathway) were visualized using geom_bar() and geom_point() ggplot2 functions.

### Bulk RNA-seq data analysis

RNA used for bulk RNA sequencing were extracted from RGCs isolated from the injured eye at 2 days post-ONC (n = 3 eyes/group, 1 eye/mouse). Raw data (raw reads) of .fastq format were first processed through in-house perl scripts by Novogene. In this step, clean data (clean reads) were obtained by removing reads containing adapter, ploy-N and low-quality reads from raw data. At the same time, Q20, Q30 and GC content of the clean data were calculated. All the downstream analyses were based on clean data with high quality.

Reference genome and gene model annotation files were downloaded from the genome website directly. The index of the reference genome was built using Hisat2 v2.0.5 and paired-end clean reads were aligned to the reference genome using Hisat2 v2.0.5^37^. The mapped reads of each sample were assembled by StringTie (v1.3.3b)^38^ in a reference-based approach.

The reads numbers mapped to each gene was counted by featureCounts v1.5.0-p3^39^. FPKM of each gene was calculated based on the length of the gene and reads count mapped to this gene. Differential expression analysis of 3 conditions/groups (3 biological replicates per condition) was performed using the DESeq2 R package (1.20.0)^40^. DESeq2 provides statistical routines for determining differential expression in digital gene expression data using a model based on the negative binomial distribution. The resulting P-values were adjusted using Benjamini and Hochberg’s approach for controlling the false discovery rate. Genes with an adjusted P-value ≤ 0.05 found by DESeq2 were assigned as differentially expressed (DEGs).

The KEGG and GO pathway annotations of the DEGs were performed with ShinyGo^41^ version 0.76 with the FDR cutoff = 0.05. The top 10 up or down-regulated pathways that are relevant to RGCs were visualized in dotplots (input genes ≥ 5 per pathway, sorted by fold enrichment). For GSEA pathway analysis of the DESeq2 output, we utilized clusterProfiler^42^, msigdbr^43^, and org.Mm.eg.db^44^. For the pathways, Hallmark category (category = “H”)^45^ was used, and upon filtering and sorting data based on log2FC values, we generated the table with GSEA() function. The top RGC-relevant pathways were visualized using ggplot2 package (setSize ≥ 5, sorted by Normalized enrichment score, NES).

The volcano plot was generated with the DESeq2 output using the Enhanced Volcano package^46^. The genes labeled on the plot are the result of two approaches: 1) the most significant genes appearing in DESeq2 output by log2FC and adjusted p-value; 2) genes appearing significant in the single nuclei RNA-seq data, pathways related to differentially methylated regions, and experimental data.

### Cell-cell interactions atlas generation

Upon generating the CellChat objects, we exported the indegree and outdegree representing the incoming and outgoing signaling, respectively. Those were used to generate the matrix consisting of cell barcodes as row names, while pathways (incoming), pathways (outgoing), ligands – ligand-receptor pairs:pathways (outgoing), receptors – ligand-receptor pairs:pathways (incoming) were used as column names. The CellChat object exported values were used to fill in the matrix. The following objects generated were formatted to follow the unified structure of the objects to be deposited to the Broad Institute’s Single Cell Portal.

## Data Availability

The raw and processed data of bisulfite, bulk, and single nuclei transcriptomic experiments generated in this study have been deposited in the GEO database under accession codes: <u>GSE229033</u> (bulk), <u>GSE229034</u> (bisulfite), <u>GSE228627</u> (single-nuclei). The single nuclei data can be explored at the Broad Institute’s Single Cell Portal: https://singlecell.broadinstitute.org/single_cell/study/SCP2321/. The cell-cell interactions can be explored at: https://singlecell.broadinstitute.org/single_cell/study/SCP2333/ for the Control, and https://singlecell.broadinstitute.org/single_cell/study/SCP2351/ for the *CKO^Dnmt3a+/-^*. The toolbox used to perform the data analysis is described in **Supplementary Table 3**.

## Code Availability

The code for reproducing the bioinformatical analysis is available on the following GitHub repository: https://github.com/mcrewcow/Lydia_ChenLab_RGC_pONC_retina.

## Intravitreal injection in mice

Procedure for intravitreal injection was essentially described^6^. During intravitreal injection, a pulled microcapillary tube was inserted just behind the ora serrata in the mouse peripheral retina. The vitreous was then partially removed (about 2 µl) to enable the injection of 2 µl of solution into the vitreous chamber. For experiments using AAVs, mice were kept in housing for at least two weeks after the injection of AAV to ensure that the target genes were stably expressed. For tracing regenerating axons anterogradely, 2 µl of 2 µg/µl CTB (List Biological Laboratories, 104) was injected intravitreally 2-3 days before mice were sacrificed at various time points to assess the extent of nerve regeneration after ONC.

## Adeno-associated viruses (AAV) constructs and production

AAV serotype 2/2 (AAV2) was used in all virus-mediated knockdown experiments. For Dnmt3a gene knockdown, vectors were generated in-house where an AAV.Scramble (scrambled control (5’ - cctaaggttaagtcgccctcgctcgagcgagggcgacttaaccttagg – 3’) and two AAV.shRNAs targeting Dnmt3a (#1: 5’ - cgctccgctgaaggaatatttctcgagaaatattccttcagcggagcg – 3’ and #2: 5’ - ggcatccactgtgaatgataactcgagttatcattcacagtggatgcc – 3’) were designed using the GPP Web Portal (Broad Institute). The shRNA sequences were synthesized as single-stranded oligonucleotides, annealed, and inserted into pAAV-U6-sgRNA-CMV-GFP (Addgene #85451) at SapI and SapI sites. AAV2 were produced as described using a triple transfection approached in HEK293T cells in 15 cm dishes with polyethylenimine and were harvested after 60 h incubation, purified by iodixanol gradient ultracentrifugation, and quantified by quantitative PCR (titers: >1 × 10^12^ genome copies per ml). AAV2/2.CASI.CRE.RBG (AAV.Cre) and AAV2/2.CASI.EGFP.RBG (AAV.GFP) (titers: >1.02 × 10^12^ GC/mL) were purchased from the Gene Transfer Vector Core at Schepens Eye Research Institute.

