## Extended Figures with Legends for "Suppressing DNMT3a Alleviates the Intrinsic Epigenetic Barrier for Optic Nerve Regeneration and Restores Vision in Adult Mice"

### Extended Data Fig.1

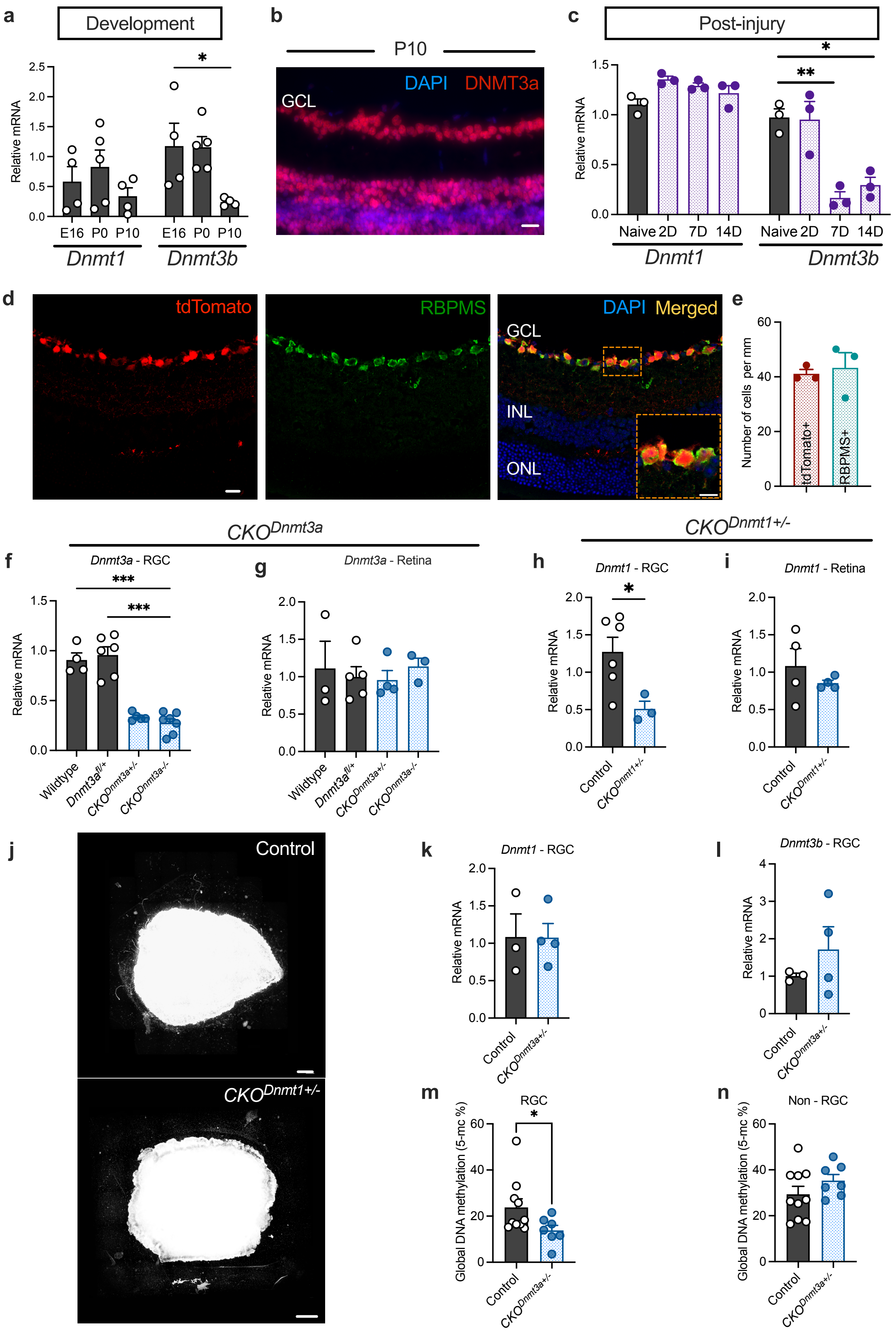

**Extended data Fig. 1. DNMT expression during development or injury in RGCs of wildtype and mutant mice. a,** qPCR quantification of *Dnmt1* and *Dnmt3b* expression in RGCs isolated from E16, P0 and P10 mice. **b**, Representative image of retinal sections taken from a P10 mouse that was immunolabeled for DNMT3a (red) and counter-stained with DAPI (blue), showing intensive DNMT3a signal in the ganglion cell layer (GCL) and other retinal layers. Scale bar: 20 µm. **c**, qPCR quantification of *Dnmt1* and *Dnmt3b* expression in RGCs taken at day 2 to 14 after ONC. **d**, Representative images of retinal sections taken from a *Vglut2-Cre:R26-tdTomato* mouse that were immunolabeled for RBPMS (green) and counter-stained with DAPI, showing colocalization (orange) of tdTomato red (*Cre*+) and RBPMS+ cells. INL, inner nuclear layer; ONL, outer nuclear layer. Scale bar: 20 µm; inset: 10 µm. **e**, Counts of tdTomato red (*Cre*+) and RBPMS+ cells in *Vglut2-Cre:R26-tdTomato* mouse retinal sections. **f-g**, Gene expressions of *Dnmt3a* in the RGCs (**f**) and retinas (**g**) of wildtype, mutant, and corresponding littermate *fl/+* control mice. **h-i**, Gene expression of *Dnmt1* in the RGCs (**h**) and retinas (**i**) of $CKO^{Dnmt1+/-}$ and littermate control mice. **j**, Representative images of cultured retinal explants derived from control (left) and $CKO^{Dnmt1+/-}$ mice immunolabeled for β-III tubulin. **k, l,** qPCR quantification of *Dnmt1* (**k**) and *Dnmt3b* (**l**) expression in RGCs of $CKO^{Dnmt3a+/-}$ mice. **m, n**, Levels of global DNA methylation (5mc%) detected in RGCs and non-RGC retinal cells of control and $CKO^{Dnmt3a+/-}$ mice (n ≥ 3 mice/group; $*P < 0.05$, $**P < 0.01$, $***P < 0.001$; for **a**, **c**, **f**, **g**, one-way ANOVA; for **e**, **h**, **i**, **k-n**, unpaired t-test; mean ± s.e.m.).

Extended Data Fig. 2

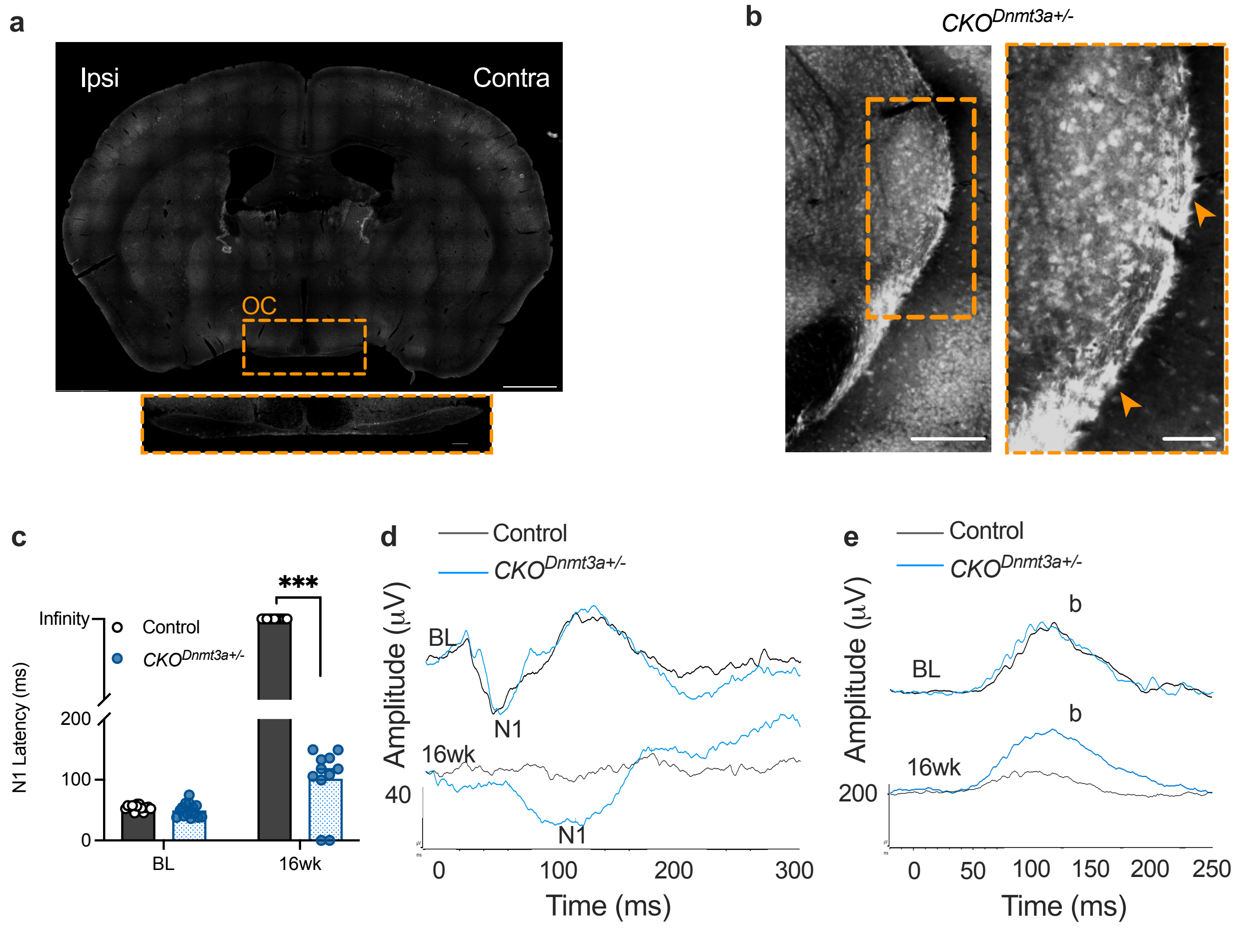


**Extended Data Fig. 2. Reinnervation of brain target by regenerated axons and functional recovery following ONC in *CKO*$^{Dnmt3+/-}$ mice. a,** Representative image of brain section taken from a control mouse post-ONC showing absence of CTB-labeling in the optic chiasm (OC). Ipsi, ipsilateral; Contra, contralateral. Scale bar: 1 mm, inset: 100 µm. **b,** Image of brain section taken at the ventral LGN (vLGN) level of a *CKO*$^{Dnmt3+/-}$ mouse at 16 weeks post-ONC, showing CTB-labeled axons entering the contralateral vLGN. Scale bar: 200 µm; inset: 100 µm. Insets are circled by orange dash lines. Arrows denote the vLGN. **c**, VEP N1 latency measured in *CKO*$^{Dnmt3a+/-}$ and littermate control mice before (BL) and at 16 weeks post-ONC. Note the prolonged N1 latency in optic nerve-injured *CKO*$^{Dnmt3a+/-}$ mice compared to BL while N1 waveform was not detected in control mice, so the N1 latency is expressed as indefinite. ($n \geq 11$ mice/group; $^{***}P < 0.001$, multiple unpaired t-tests; mean ± s.e.m.). **d, e**, Representative waveforms of the VEP (**d**) and pSTR (**e**) taken from *CKO*$^{Dnmt3a+/-}$ and littermate control mice before (BL) and at 16 weeks post ONC.

**Extended Data Fig. 3**

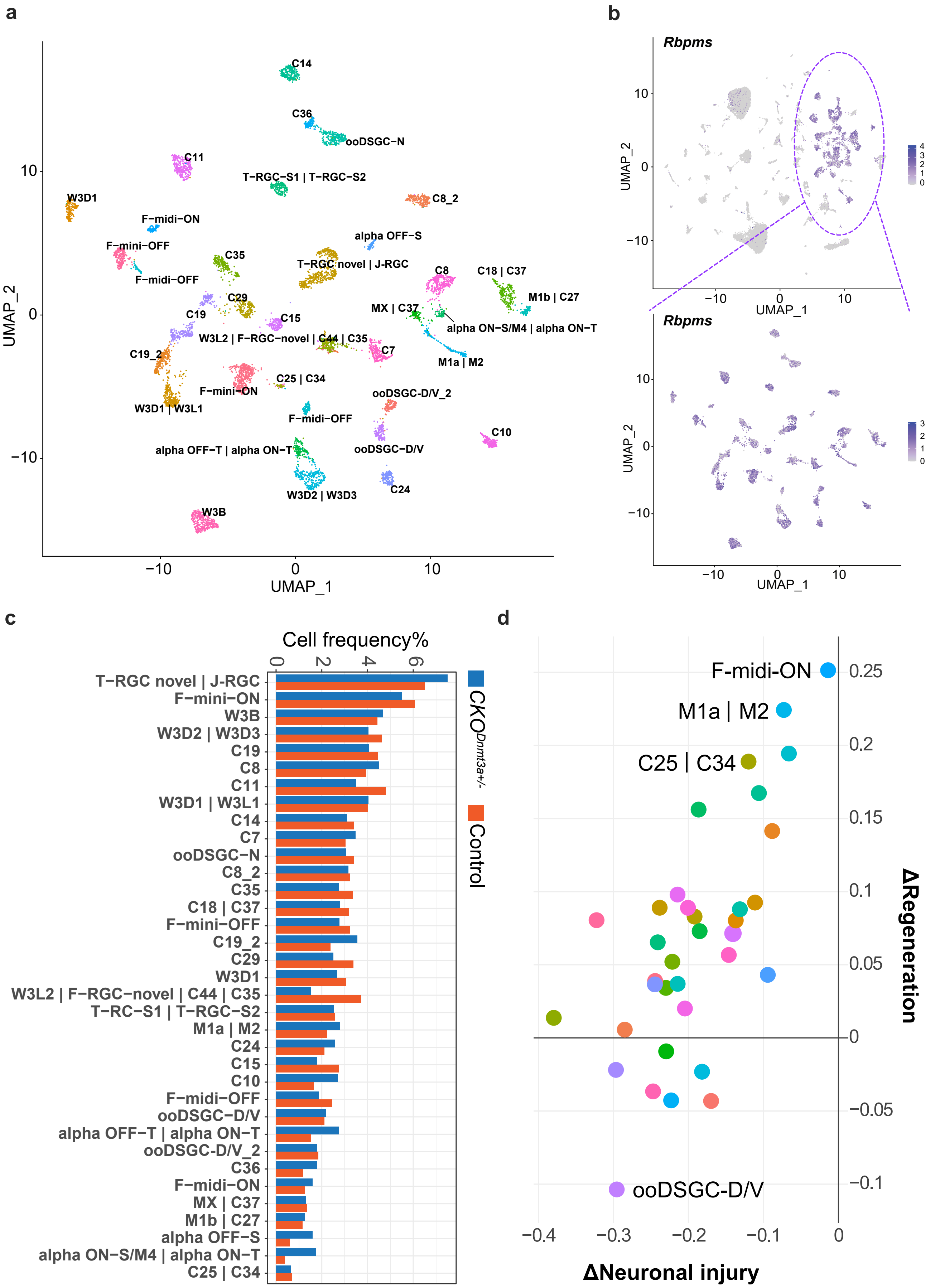

**Extended Data Fig. 3. Alterations in the RGC transcriptomes but not type distributions. a**, Dimplot showing RGC clustering in snRNA-seq dataset merged from control and $CKO^{Dnmt3a+/-}$ mice and identifying 35 clusters with 42 out of 45 RGC types (annotation showing multiple RGC types, e.g. W3D2 | W3D3, in some clusters). **b,** Feature plot showing expression of *Rbpms*, a pan-RGC marker, in the entire retinal cell clusters and RGC subsets in merged snRNA-seq datasets of control and $CKO^{Dnmt3a+/-}$ mice. **c**, Barplot showing RGC types frequency with the total number of RGCs to be 3791 for control and 3319 for $CKO^{Dnmt3+/-}$, respectively. **d**, Scatterplot visualization of relative strength of combined Gene set enrichment analysis (GSEA) pathway scores related to neuronal injury (n = 4 GSEA pathways) and nerve regeneration (n = 2 GSEA pathways) between $CKO^{Dnmt3+/-}$ and control RGC types. The difference was calculated by subtraction of the normalized average expression in $CKO^{Dnmt3+/-}$ RGCs from the Controls. Each colored dot represents a RGC type noted in the dimplot (**a**). A 3D visualization of this scatterplot is in the Extended Data HTML File.

**Extended Data Fig. 4**

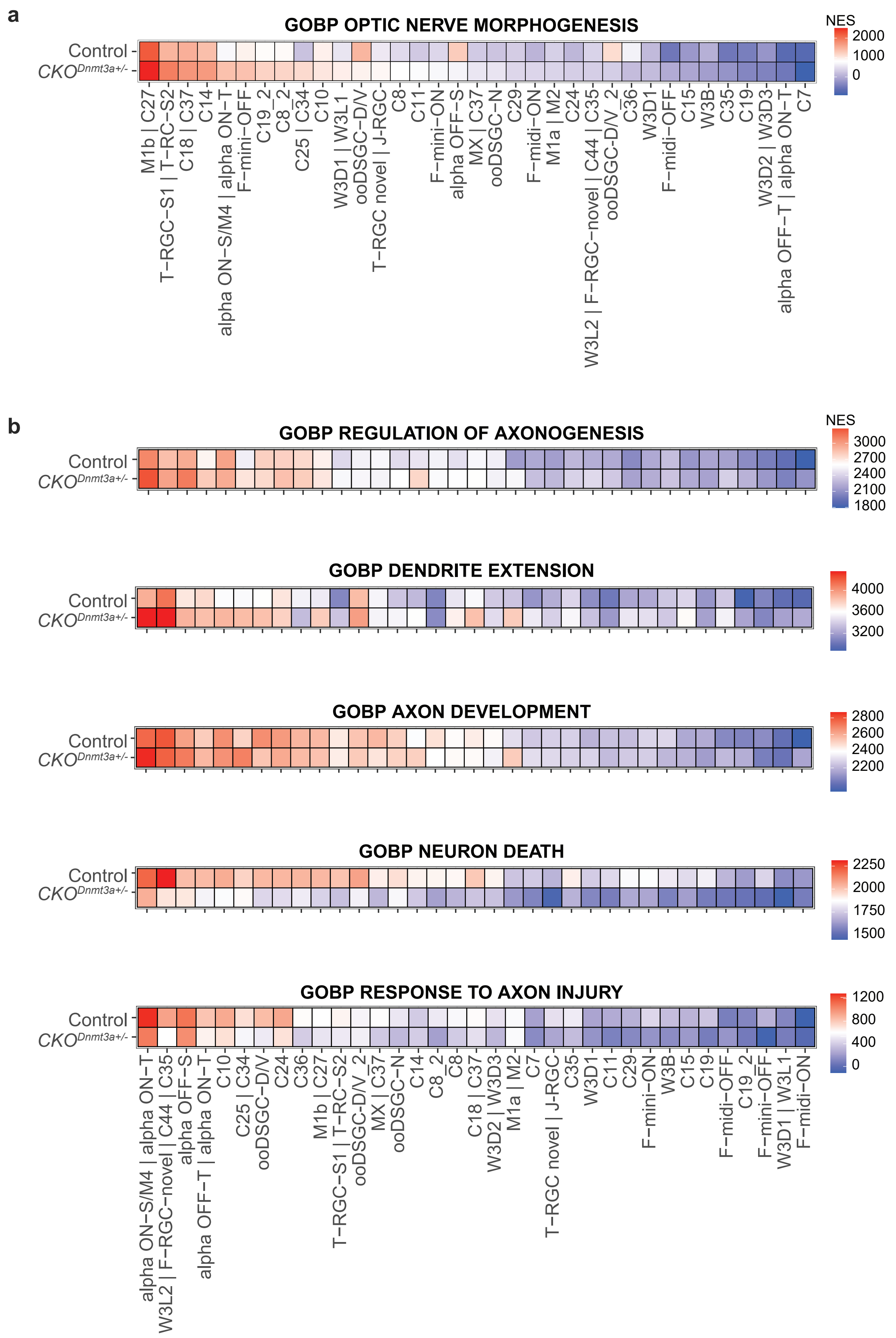

**Extended Data Fig. 4. Wide-scale shifts of transcriptome profiles in RGC types. a-b,** Heatmaps of Gene ontology biological processes (GOBP) analysis for snRNA-seq DEGs in individual RGC clusters indicating increased strengths in optic nerve morphogenesis (**a**), axonogenesis, dendritic extension, and axon development (**b**), but decreased strengths in neuronal death and response to axon injury (**b**). NES, normalized enrichment scores.

### Extended Data Fig. 5

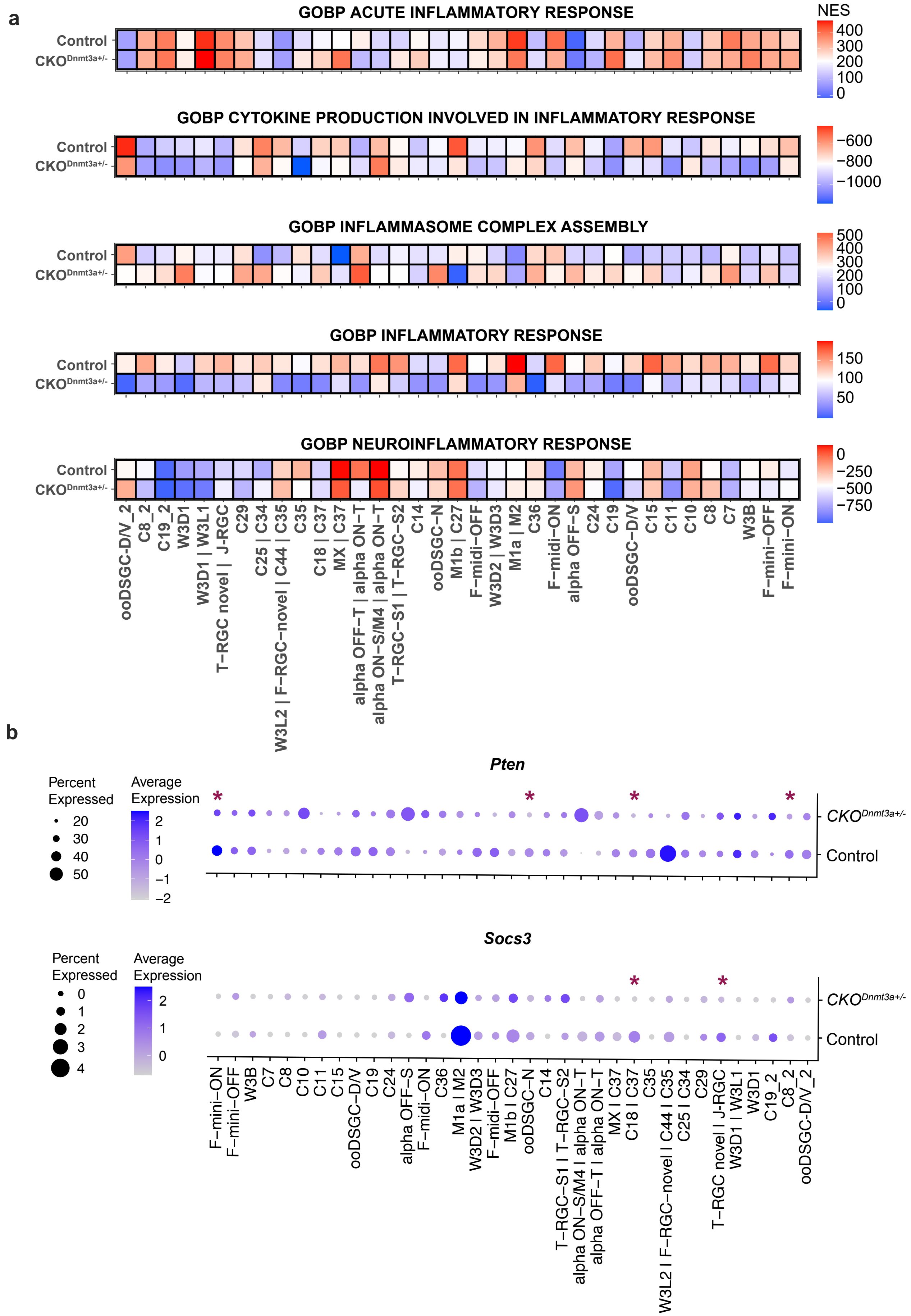

**Extended Data Fig. 5. Comparison of *Pten* and *Socs3* expression in various RGC types of *CKO*$^{Dnmt3+/-}$ and control mice. a,** Heatmaps of Gene ontology biological processes (GOBP) analysis for snRNA-seq DEGs in individual RGC clusters indicating reduced acute inflammatory response, and inflammatory cytokine production in *CKO*$^{Dnmt3+/-}$ RGCs compared to controls. NES, normalized enrichment scores. **b**, Dotplot visualization of *Pten* (top) and *Socs3* (bottom) expression in individual RGC clusters detected by snRNA-seq analysis of *CKO*$^{Dnmt3+/-}$ and control RGCs. Red stars denote RGC types with significantly downregulated expressions of *Pten* or *Socs3* in the *CKO*$^{Dnmt3+/-}$ compared to the control ($P < 0.05$).

### Extended Data Fig. 6

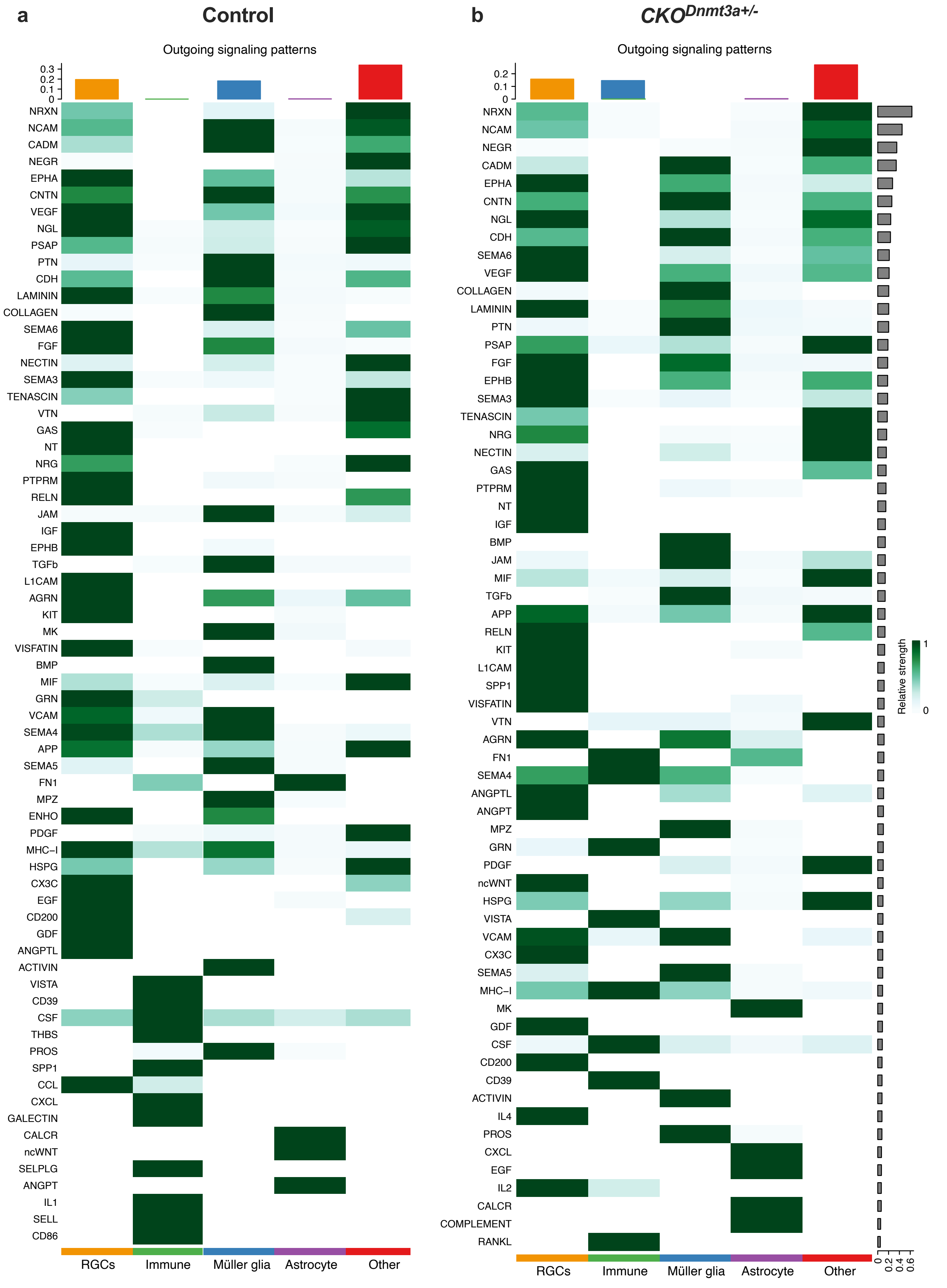

**Extended Data Fig. 6. CellChat predictions of outgoing signals from different retinal cell types. a**, **b**, CellChat heatmap representing all predicted outgoing signaling patterns from retinal cells of control (**a**) and $CKO^{Dnmt3+/-}$ (**b**) mice. The cell types analyzed include RGC, immune cells (including microglia/macrophages, T helpers and T cells), Müller glia, amacrine, and other (including horizontal cells, bipolar neurons, astrocytes, retinal pigment epithelium, rods, and cones). The top colored bar plot demonstrates the total signaling strength of a cell group by summarizing all signaling pathways shown on the heatmap. X-axis denotes the cell populations, and Y-axis denotes the pathways in an order of signaling strength/contribution.

### Extended Data Fig. 7

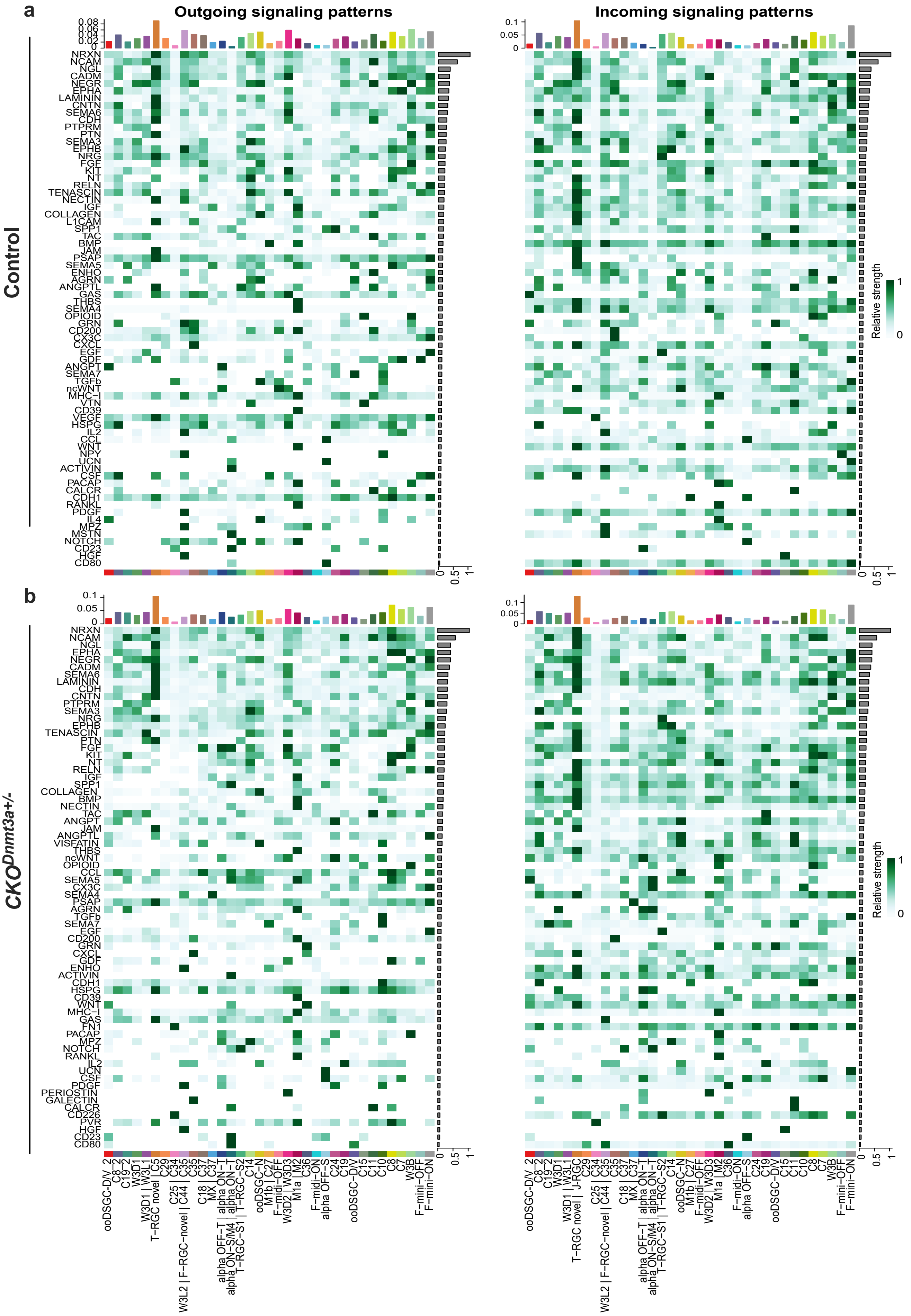

**Extended Data Fig. 7. CellChat predictions of incoming and outgoing signals sent within RGC types. a, b,** CellChat heatmap representing all predicted incoming and outgoing signaling patterns among RGC types of control (**a**) and $CKO^{Dnmt3+/-}$ (**b**) mice. Colored bars shown on top of the heatmap illustrate the total signaling strength of a cell group by summarizing all signaling pathways. X-axis denotes the RGC types, and Y-axis denotes the pathways in an order of signaling strength/contribution.

**Extended Data Fig. 8**

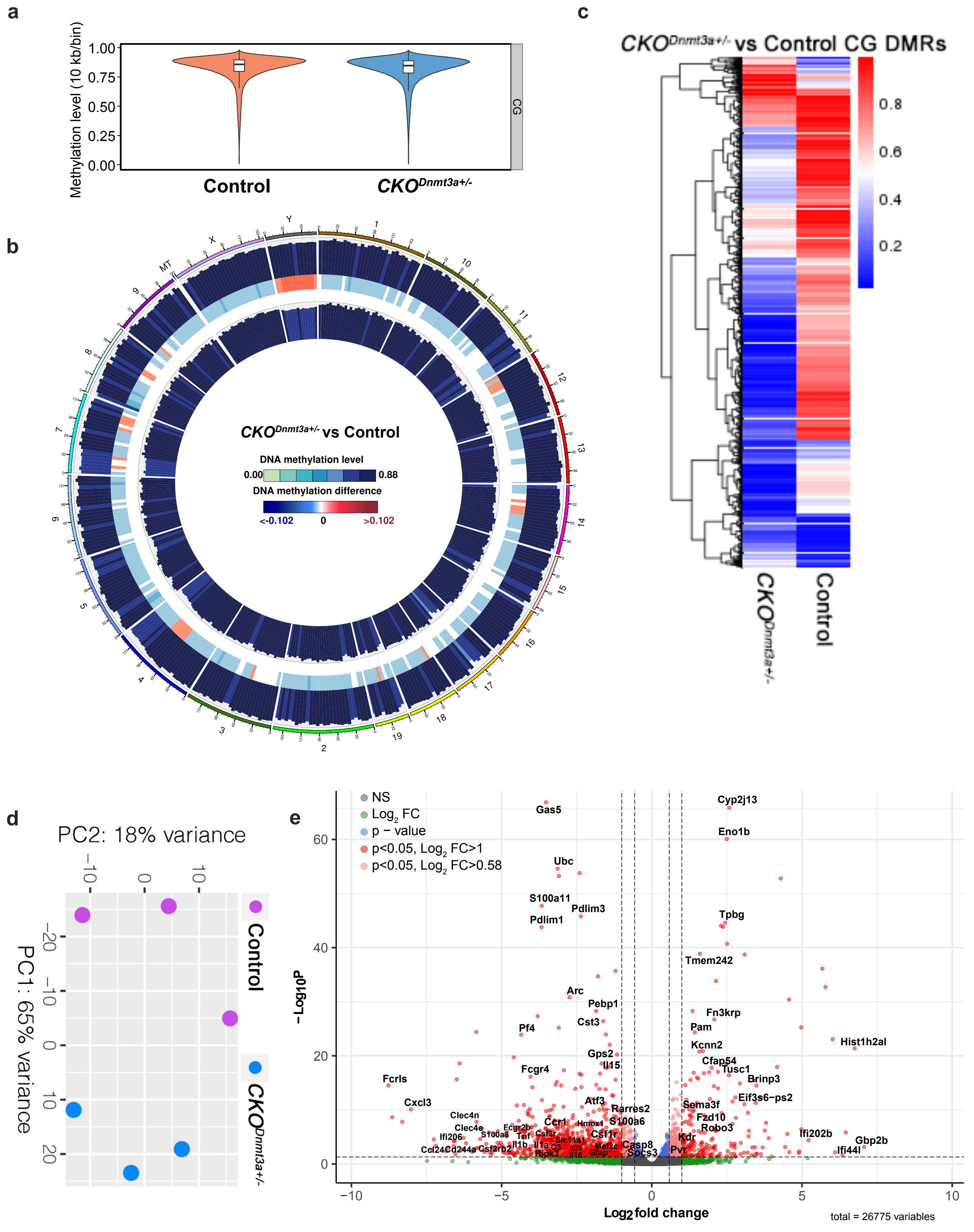

**Extended Data Fig. 8. DMR demethylation and upregulation of axon regeneration pathways by *Dnmt3a* deficiency.** **a**, The total methylation level of CG in the RGCs of control and $CKO^{Dnmt3a+/-}$ mice at 2-day post-crush (n = 2 mice/group). **b**, Circos plot of CG methylation level and the difference between control and $CKO^{Dnmt3a+/-}$ RGCs. From outside to inside, each ring represents: 1. methylation level for the $CKO^{Dnmt3a+/-}$ group, 2. methylation level difference between control and $CKO^{Dnmt3a+/-}$ (heatmap) 3. methylation level for the control group. The chromosomes are divided into bins where methylation level of each bin is calculated as the number of reads with methylation / (number of reads with methylation + number of reads without methylation). **c**, Cluster heatmap for CG methylation level at the differentially methylated regions (DMRs). **d**, Principle component analysis of bulk RNA-seq data (n = 3 mice/group), demonstrating the variability of the samples studied. **e**, Volcano plot of DEGs between $CKO^{Dnmt3+/-}$ and control RGCs by bulk RNA-seq. NS, non-significant; $Log_2$ FC, DEGs with $log_2$ fold change > 1 and p-value > 0.05; p - value, DEGs with p - value < 0.05 and $log_2$ fold change < 1.

### Extended Data Fig. 9

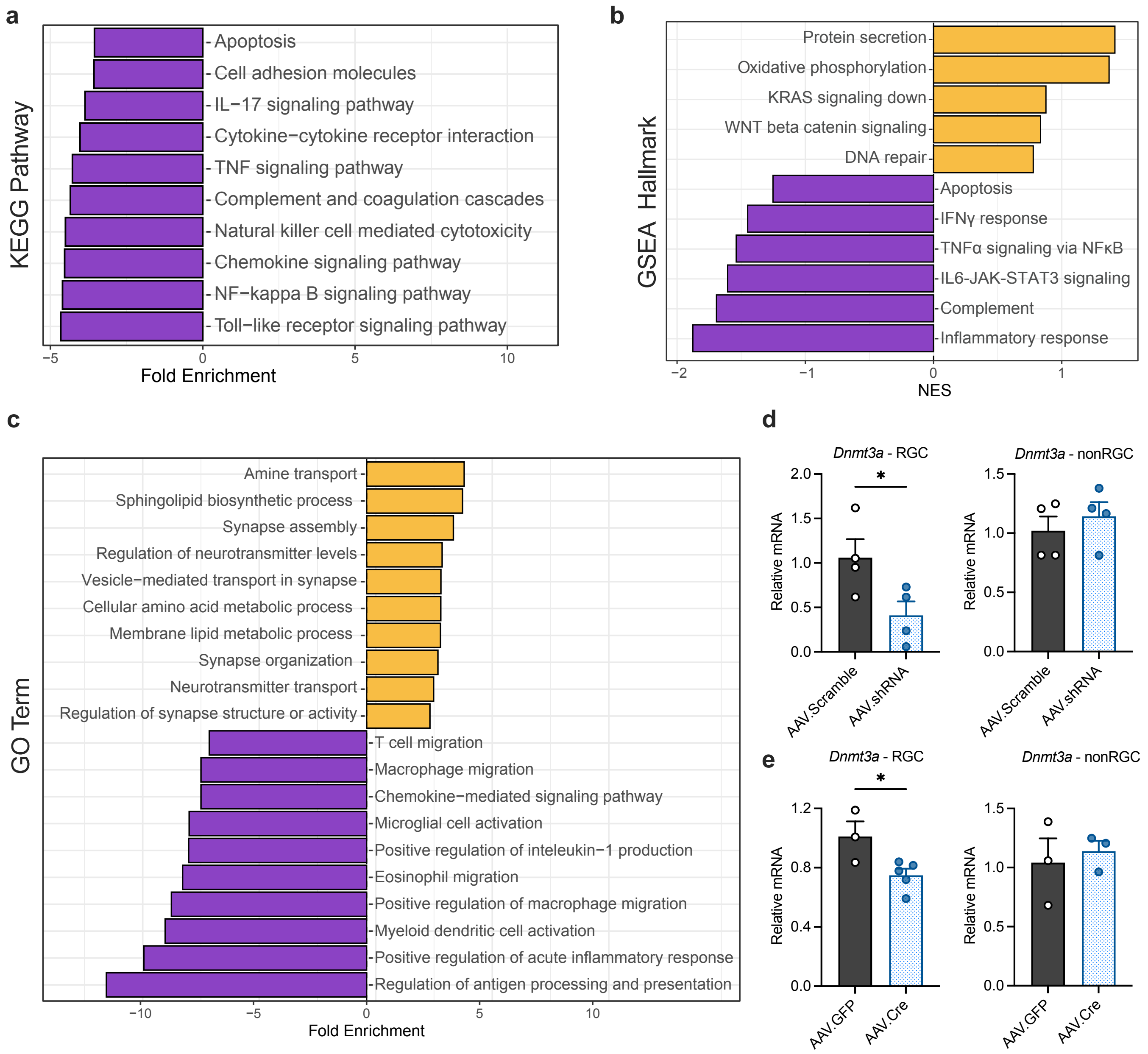

**Extended Data Fig. 9. Knockdown of *Dnmt3a* by AAV-mediated gene delivery. a**, Kyoto encyclopedia of genes and genomes (KEGG) pathways enriched by downregulated genes in bulk RNA-seq. **b**, Gene set enrichment analysis (GSEA) Hallmark pathways and **c**, Gene ontology (GO) Terms enriched by upregulated (labeled in yellow) and downregulated (labeled in purple) genes in bulk RNA-seq. **d**, qPCR quantification of *Dnmt3a* expression in RGCs and non-RGC retinal cells in adult wildtype mice at 14 days after receiving intravitreal injection of *AAV.Scramble* or *AAV.shRNA* that specifically targets *Dnmt3a*. **e**, qPCR quantification of *Dnmt3a* expression in RGCs and non-RGC retinal cells in adult *Dnmt3a* $^{fl/+}$ mice at 14 days after receiving intravitreal injection of *AAV.GPF* or *AAV.Cre* ($n \geq 3$ mice/group; **P* < 0.05, unpaired t-test; mean ± s.e.m.).

**Extended Data HTML File**

Interactive 3D scatterplot showing the distribution of RGC types in 3 dimensions that denote regeneration, injury, and inflammation pathways, respectively. The values on the axis were obtained by subtracting the normalized expression strength of specific pathways calculated in individual RGC types of *CKO*$^{Dnmt3+/-}$ mice from the corresponding RGC types of the control mice. The data showed that all RGC types/clusters revealed decreased expression in the injury-related pathways upon *Dnmt3a* deficiency, and 24 out of 35 RGC clusters exhibited increased regeneration signalings.
